# Modulation of the cAMP levels with a conserved actinobacteria phosphodiesterase enzyme reduces antimicrobial tolerance in mycobacteria

**DOI:** 10.1101/2020.08.26.267864

**Authors:** Michael Thomson, Kanokkan Nunta, Ashleigh Cheyne, Yi liu, Acely Garza-Garcia, Gerald Larrouy-Maumus

## Abstract

Antimicrobial tolerance (AMT) is the gateway to the development of antimicrobial resistance (AMR) and is therefore a major issue that needs to be addressed.

The second messenger cyclic-AMP (cAMP), which is conserved across all taxa, is involved in propagating signals from environmental stimuli and converting these into a response. In bacteria, such as *M. tuberculosis*, *P. aeruginosa*, *V. cholerae* and *B. pertussis*, cAMP has been implicated in virulence, metabolic regulation and gene expression. However, cAMP signalling in mycobacteria is particularly complex due to the redundancy of adenylate cyclases, which are enzymes that catalyse the formation of cAMP from ATP, and the poor activity of the only known phosphodiesterase (PDE) enzyme, which degrades cAMP into 5’- AMP.

Based on these two features, the modulation of this system with the aim of investigating cAMP signalling and its involvement in AMT in mycobacteria id difficult.

To address this pressing need, we identified a new cAMP-degrading phosphodiesterase enzyme (Rv1339) and used it to significantly decrease the intrabacterial levels of cAMP in mycobacteria. This analysis revealed that this enzyme increased the antimicrobial susceptibility of *M. smegmatis* mc^2^155. Using a combination of metabolomics, RNA-sequencing, antimicrobial susceptibility assays and bioenergetics analysis, we were able to characterize the molecular mechanism underlying this increased susceptibility.

This work represents an important milestone showing that the targeting of cAMP signalling is a promising new avenue for antimicrobial development and expands our understanding of cAMP signalling in mycobacteria.

## INTRODUCTION

Along with climate change and viral pandemics, antimicrobial resistance (AMR) is currently one of the greatest threats to human health. Antimicrobial tolerance (AMT) precedes antimicrobial resistance^1–3^, and although AMR stems from heritable genetic changes in bacteria, such as the expression of efflux pumps or antibiotic degrading enzymes, AMT is mediated by transient and reversible physiological adaptations in signalling systems that orchestrate the remodelling of bacterial physiology to alter their susceptibility to antimicrobials.

Cyclic AMP signalling, which is one of the best studied signalling pathways in biology, has been implicated in AMR^4,5^. Several earlier studies have linked cAMP signalling to antibiotic resistance in gram-negative bacteria^6,7^. For example, Alper *et al*. demonstrated that *Salmonella typhimurium* with mutations in the cAMP control system, which comprises adenylyl cyclase and the cAMP receptor protein, lead to a partial resistance to 20 commonly used antibiotics^6^. Moreover, Kary *et al*. reported that *Salmonella typhimurium* strains lacking Crp (a cAMP-activated global transcriptional regulator) or with altered levels of cAMP exhibit reduced susceptibility to fluoroquinolones as a result of decreased permeability and increased efflux of this class of antibiotics^7^.

Despite the above-mentioned research on gram-negative pathogens, the link between cAMP signalling and AMR/AMT in mycobacteria, including the strict human pathogen *Mycobacterium tuberculosis*, the vaccine strain *Mycobacterium bovis* BCG and the model organism *Mycobacterium smegmatis*, has not been well studied. This lack of knowledge can be explained by the complexity of the cAMP signalling system in mycobacteria and the lack of effective tools for manipulating this system^8–12^. In contrast to the model organism *E. coli*, which comprises one enzyme that generates cAMP, adenylate cyclase (Cya), and one enzyme that hydrolyses cAMP, a phosphodiesterase (PDE) (CpdA)^13^. Conversely, the *M. tuberculosis* H37Rv genome encodes 16 adenylate cyclases belonging to the class-III adenylate cyclase enzymes, which typically harbour a cAMP-producing domain linked to a sensory domain^12,14^. Interestingly, only one PDE (denoted Rv0805) found in mycobacteria belonging to the Mtb complex^15–19^, *M. leprae*, *M. marinum* and *M. ulcerans*, has been characterized even though cAMP PDE activity has also been reported in the model organism *M. smegmatis^20^*, which suggesting the presence of an additional PDE that is likely ubiquitous in the mycobacteria genus. Furthermore, Rv0805 exhibits more activity towards 2’3’-cAMP, which arises from RNA degradation, than towards the actual second messenger^19^. The high redundancy of cAMP-producing enzymes and the poor activity of the known PDE make it difficult to modulate this system with the aim of investigating the involvement of cAMP signalling with AMR/AMT in mycobacteria.

To tackle this challenge, we first searched for a tool that would allow us to efficiently modulate intracellular cAMP levels. The current exogenous strategies for altering cAMP levels, including the use of adenylate cyclase activator (forskolin) and analogues such as dibutyryl-cAMP, only result in modest changes in the intracellular cAMP levels^21,22^. Instead, we focused on finding the “missing” cAMP PDE using a combination of conventional biochemical approaches and mass spectrometry techniques (proteomics and metabolomics). With this approach, we identified Rv1339, an uncharacterized cAMP phosphodiesterase that is present in all mycobacteria. Phylogenetic analyses showed that Rv1339 is a member of a poorly characterized group of PDE enzymes with a metallo-β-lactamase fold that is present in several bacterial phyla. This group is different but evolutionarily related to the typical class-II PDEs.

To demonstrate that Rv1339 can be used as a tool to alter the intracellular cAMP pool in mycobacteria and evaluate its impact on physiology, we used the model organism *M. smegmatis* mc^2^155. The results showed that the expression of Rv1339 leads to a 3-fold decrease in the intracellular cAMP levels and thus indicate that Rv1339 likely plays a role in cAMP signalling in mycobacteria.

By examining the bacterial transcriptome, metabolome, bioenergetics and permeability, we characterized the mechanisms through which Rv1339 expression results in increased permeability and decreased AMT to two common antibiotics with different modes of action, ethambutol and streptomycin. These findings suggest that cAMP signalling might be a promising new target for the development of compounds that inhibit the AMT mechanisms of critical bacterial pathogens.

## RESULTS

### Rv1339 is an atypical class-II PDE

To identify novel sources of cAMP PDE activity in mycobacteria, we first utilized an unbiased biochemical approach involving the sequential fractionation of *M. bovis* BCG ∆Rv0805 lysate coupled to a cAMP PDE activity assay^20,23^. The enrichment of cAMP PDE activity was achieved through two chromatography steps that separate proteins based on their charge and molecular weight (Supplementary Fig. 1). After each purification step, the active factions were identified through the cAMP PDE assay and pooled for further purification. Subsequently, trypsin digestion and proteomic analysis were used to identify the proteins present in the final set of active fractions. Out of eight proteins that were uniquely present in the active fractions, only three were uncharacterized proteins: Rv0250, Rv2568 and Rv1339. Himar-1 transposon mutagenesis studies have revealed that the uncharacterized proteins Rv0250 and Rv2568 are nonessential^24^. Rv1339 is a conserved hypothetical protein that is ubiquitous in mycobacteria and is predicted to have hydrolase activity. Based on these observations, we decided to focus on the characterization of Rv1339.

Literature mining and phylogenetic analyses revealed that Rv1339, YfhI from *Bacillus subtilis^25^* and CpdA from *Corynebacterium glutamicum^26^*, are representatives of a previously unrecognized class of PDEs defined by the signature metal-binding motif [T/S]HXHXDH, where X tends to be a hydrophobic, small or very small residue. This motif is reminiscent of the bona fide class-II PDE motif HXHLDH^27^, and indeed, both families are evolutionarily related and belong to the metallo-hydrolase/oxidoreductase superfamily (SSF56281). We propose to name this novel group atypical class-II PDEs. Interestingly, atypical class-II PDEs are closely related and might have evolved from the ribonuclease Z family of proteins involved in the maturation of tRNA (Fig. 1a).

**Figure 1:**
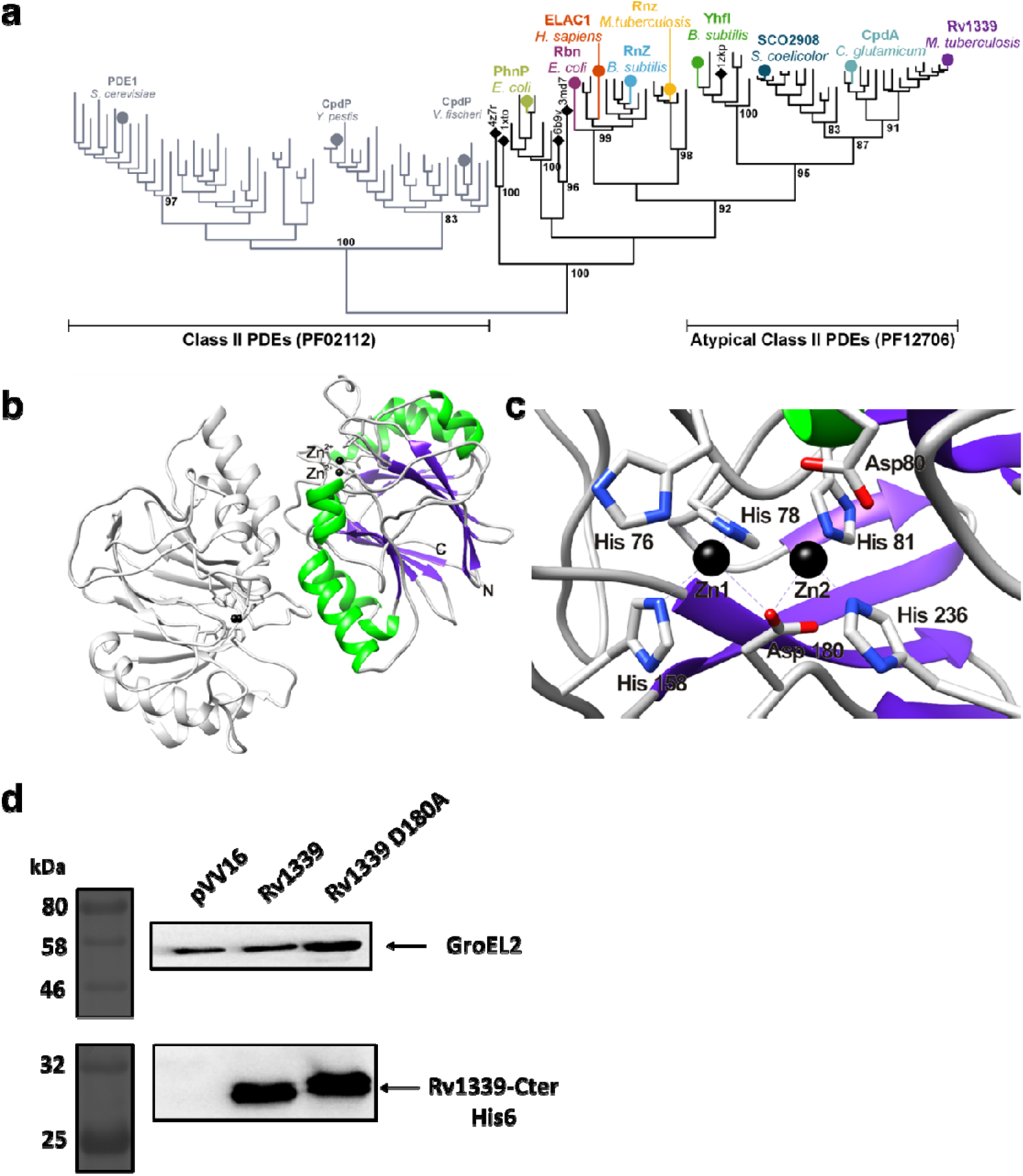
Phylogenetic analysis and expression of Rv1339 in *M. smegmatis* mc^2^155. **a**. Maximum-likelihood phylogenetic consensus tree of selected members of typical (PF02112) and atypical (PF12706) phosphodiesterase (PDE) class-II families. **b**. Ribbon representation of a homology model of Rv1339 established using Modeller 9.23^104^ with the crystal structure of the metallo-beta-lactamase fold protein YhfI from *Bacillus subtilis* (PDB: 6KNS) as the template. YhfI was found to be a homodimer in solution by size-exclusion chromatography^25^. The structure exhibits the ab-sandwich configuration characteristic of the metallo-β-lactamase fold, which consists of two β sheets of seven β strands each (purple) and α helices (green) capping the β sheet. The two zinc cations in the active sites are depicted in black. The figure was created using UCSF Chimaera^105^. **c**. Zoomed view of one of the active sites of the homology model of Rv1339 showing the coordinating histidine and aspartate residues. Asp180 was chosen for mutagenesis because it is expected to be involved in the coordination of both zinc cations. **d**. Western blot of empty control vector-, Rv1339- and Rv1339 D180- expressing *M. smegmatis* mc^2^155 strains. The proteins were probed with α-His and α-GroEL2 antibodies, and each lane was loaded with 50 μg of protein for normalization.

For the generation of a catalytically inactive form of Rv1339, we established a homology model of Rv1339 (Fig. 1b) to inform our site-directed mutagenesis efforts^28^. We mutated the aspartate residue at position 180 to an alanine and found that the initial residue was likely responsible for stabilizing both divalent metal ions in the metal-binding site (Fig. 1c). The aspartate residue at this position has been described in the literature as required for the activity of other metallo-β-lactamase proteins ^29^. The different constructs generated in this study were successfully expressed in the model organism *M. smegmatis* mc^2^155 (Fig. 1d).

### The expression of Rv1339 leads to a decrease in the intracellular cAMP levels, an increase in cAMP turnover and growth defects

To investigate the potential cAMP PDE activity of Rv1339, we first monitored the growth of *M. smegmatis* mc^2^155 with the empty control vector (pVV16), Rv1339 and Rv1339 D180A. The bacteria expressing Rv1339, but not Rv1339 D180A or the empty control vector, displayed a marked growth defect of approximately 30% when cultured in 7H9-defined media supplemented with glucose and glycerol carbon sources (Fig. 2a and Supplementary Table 1). This finding suggests that Rv1339 expression exerts a major effect on bacterial physiology. To determine whether the expression of Rv1339 in *M. smegmatis* mc^2^155 affects the cAMP levels, we measured the intracellular cAMP levels using a previously employed method^30^. The expression of Rv1339 led to a 3.2-fold decrease in the intracellular cAMP levels (Fig. 2b). In addition, we measured the incorporation of ^13^C through [U-^13^C_6_]-glucose and [U-^13^C_3_]-glycerol stable isotope tracing and found a 3-fold increase (Supplementary Fig. 2a). This result suggests that the Rv1339-expressing strain exhibits a higher turnover of cAMP than the empty control vector-expressing strain. Taken together, these data show that Rv1339 possesses cAMP PDE activity in *M. smegmatis* mc^2^155. Notably, this activity was abrogated by the Rv1339 D180A mutation (Fig. 1b), which suggested the successful generation of a Rv1339 activity mutant and demonstrated that the observed phenotype was due to the functional catalytic motif of Rv1339. These findings demonstrated that we had developed a powerful tool for altering the intracellular cAMP levels and investigating the impact of these changes on AMR/AMT.

**Figure 2:**
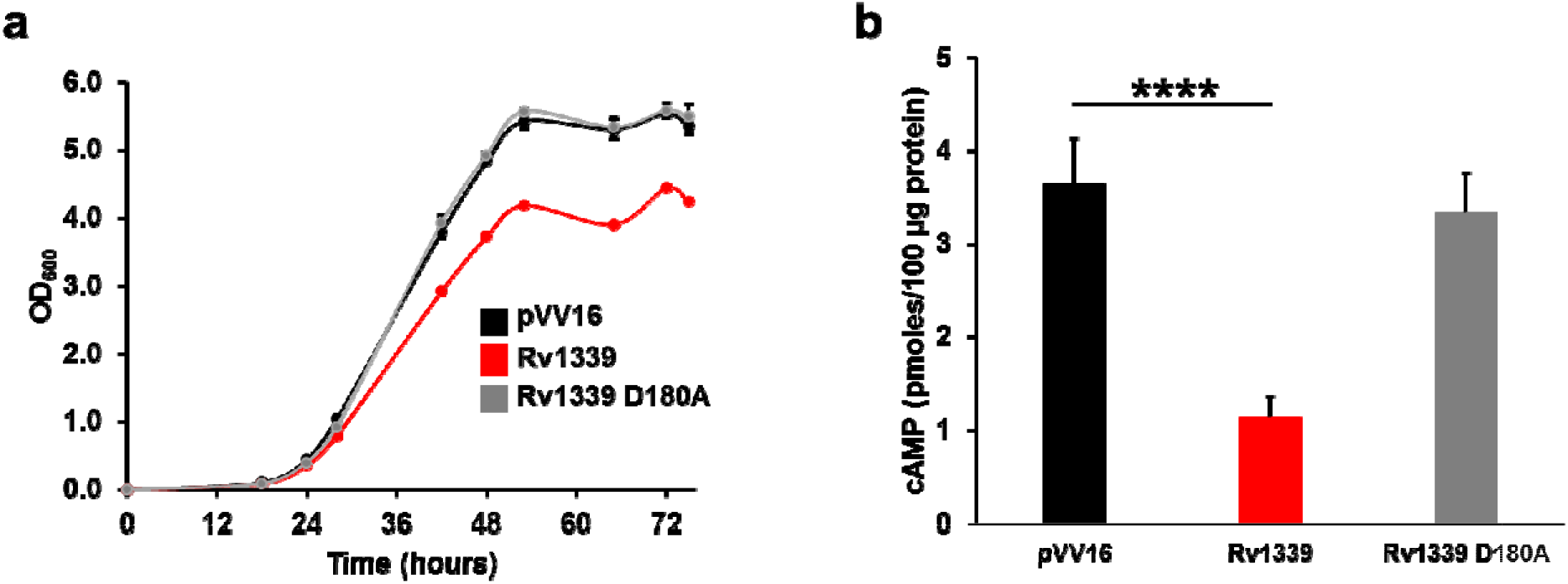
Expression of Rv1339 in *M. smegmatis* mc^2^155 alters the growth and cAMP intracellular levels. **a.** Growth curves of *M. smegmatis* mc^2^155 pVV16 (dark bar), *M. smegmatis* mc^2^155 pVV16:rv1339 (red bar) and *M. smegmatis* mc^2^155 pVV16 rv1339 D180A (grey) in conventional 7H9 broth. **b.** Intracellular cAMP levels of *M. smegmatis* mc^2^155 pVV16 (dark bar), *M. smegmatis* mc^2^155 pVV16::rv1339 (red bar) and *M. smegmatis* mc^2^155 pVV16::rv1339 D180A (grey bar). The data are presented as the means±SDs from two biological replicates and three replicates. Unpaired two-tailed Student’s t-tests were used to compare the data. *p<0.05, **p<0.01, ***p<0.001 and ****p<0.0001.

### A decrease in the intracellular cAMP levels leads to decreased antimicrobial tolerance

Increasing lines of evidence generated in recent years support the hypothesis that cAMP signalling is involved in regulating the susceptibility of bacteria to antimicrobials^7,31^. In light of our previous data, which showed that the cAMP levels can be effectively decreased (Fig. 1b), we decided to measure the susceptibility of Rv1339-expressing *M. smegmatis* mc^2^155 to streptomycin and ethambutol and compared it with that of the strain harbouring only the empty control vector. Using these two antibiotics, which have different mechanisms of action, we could investigate whether cAMP potentially plays a universal role in antibiotic tolerance. Streptomycin targets protein production by binding to the 16S subunit of the bacterial ribosome and inhibiting its effective translation^32^, whereas ethambutol targets the synthesis of arabinogalactan, which is a key mycobacterial cell wall component^33–36^.

First, we determined the minimum inhibitory concentration (MIC_90_) of both antimycobacterial drugs for *M. smegmatis* mc^2^155 expressing Rv1339 or the empty control vector. No significant differences in the MIC_90_ values were observed, although a minor change was found with streptomycin (Supplementary Table 2). This finding indicates that changes in the intracellular cAMP levels did not alter the resistance profile of the bacteria. We then investigated whether changes in the cAMP levels altered AMT by measuring the time-to-killing (TTK) of the empty vector control (solid lines) and Rv1339-expressing (dashed lines) *M. smegmatis* mc^2^155 strains in response to treatment with these antibiotics (Fig. 3). Specifically, the bacteria were treated with 0 x MIC_90_ (green lines), 1 x MIC_90_ (blue lines) or the clinically relevant *C*_max_ (peak levels of antibiotic found in patient serum: 20 x MIC_90_ for streptomycin and 5 x MIC_90_ for ethambutol^37^ – red lines), and the bacterial colony-forming units (CFUs) were quantified at various times during the treatment.

**Figure 3:**
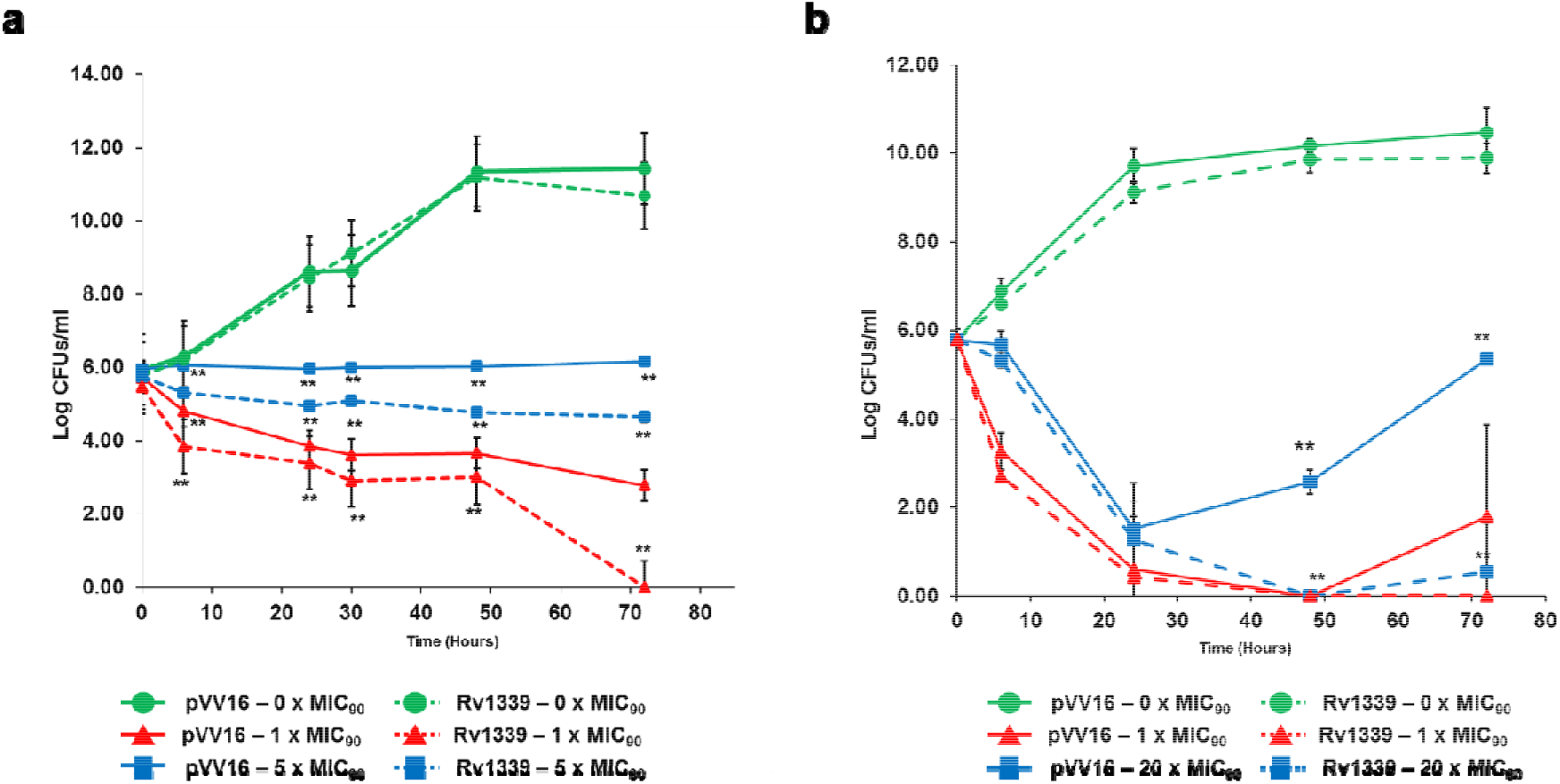
Time-to-killing assay results for the empty control vector-, Rv1339- and Rv1339 D180A-expressing *M. smegmatis* mc^2^155 strains in the presence of streptomycin (a) and ethambutol (b). The data are presented as the means±SDs from three biological replicates and three replicates. Unpaired two-tailed Student’s t-tests were used to compare the data. *p<0.05, **p<0.01, ***p<0.001 and ****p<0.0001.

The CFUs of Rv1339-expressing bacteria treated with 1 x MIC_90_ streptomycin (Fig. 2b) decreased over time from the initial 6 log_10_ to less than 2 log_10_ CFUs/ml and were considered sterilised. After 72 hours of treatment with 1 x MIC_90_ streptomycin, the bacteria harbouring the empty control vector displayed regrowth back to the baseline level at the start of the assay. This regrowth was not observed in the Rv1339-expressing bacteria, and indeed, the difference in growth between the Rv1339- and empty control vector-expressing bacteria after 48 and 72 hours of treatment at this concentration was highly significant (p<0.001). Due to the only slight difference in the MIC_90_, this finding indicates that reducing the cAMP levels abrogated the development of a tolerant population of bacteria during treatment with streptomycin at 1 x MIC_90_. In contrast, during treatment at the *C*_max_ concentration, this regrowth was not observed with either strain (Fig. 2b), which is consistent with the bactericidal nature of streptomycin.

The time-to-kill assay with the bacteriostatic antibiotic ethambutol (Fig. 2a) at 1 x MIC_90_ revealed that both the Rv1339- and empty control vector-harbouring bacteria did not display growth, which is consistent with the bacteriostatic nature of ethambutol^38^. However, Rv1339- expressing bacteria were significantly less tolerant (p<0.001), as demonstrated by a 1 x log_10_ decrease in the CFUs after 72 hours of treatment, and this finding indicated bactericidal activity of ethambutol in the presence of lower intracellular cAMP levels. Furthermore, at the *C*_max_ concentration, ethambutol exhibited modest bactericidal activity against the empty control vector-expressing strain but induced complete sterilisation of Rv1339-expressing bacteria after 72 hours of treatment.

Taken together, these data clearly indicate that a decrease in the intracellular cAMP levels renders mycobacteria less tolerant to antimicrobials. Additionally, a 3.2-fold decrease in bacterial cAMP levels was sufficient to convert a bacteriostatic antibiotic, such as ethambutol, into a bactericidal antibiotic. Because our data suggested a link between the cAMP levels and antimicrobial tolerance in *M. smegmatis* mc^2^155, we subsequently sought to investigate the mechanism underlying the decrease in tolerance observed with lower cAMP levels. Ethambutol and streptomycin have very different mechanisms of action^32,34,38^, and the finding that the Rv1339-expressing *M. smegmatis* mc^2^155 strain exhibited increased susceptibility to both antibiotics suggested that a global change in bacterial physiology occurred.

### A decrease in the intracellular cAMP levels alters the transcriptome by decreasing the expression of genes that are known to be regulated by cAMP

The best characterized mycobacterial cAMP-binding effector proteins are Crp and Cmr in *M. tuberculosis* H37Rv^13,39–42^. These transcription factors regulate various processes ranging from virulence^43–45^ to carbon metabolism^15,18,46–48^ and dormancy^39,49^. It is therefore likely that decreased cAMP levels would alter the transcriptome, and this effect could underpin the likely generalized change in the bacterial physiology that led to the increased antimicrobial susceptibility observed with the Rv1339-expressing bacteria. To investigate this hypothesis, RNA sequencing of the Rv1339- and empty control vector-expressing *M. smegmatis* mc^2^155 strains was performed, and the results were compared. Prior to the analysis, the bacteria were grown to the mid-log phase in conventional 7H9 with glycerol and glucose as the carbon sources. After RNA sequencing, the differentially expressed genes were analysed (Fig. 4 and Supplementary Table 3).

**Figure 4:**
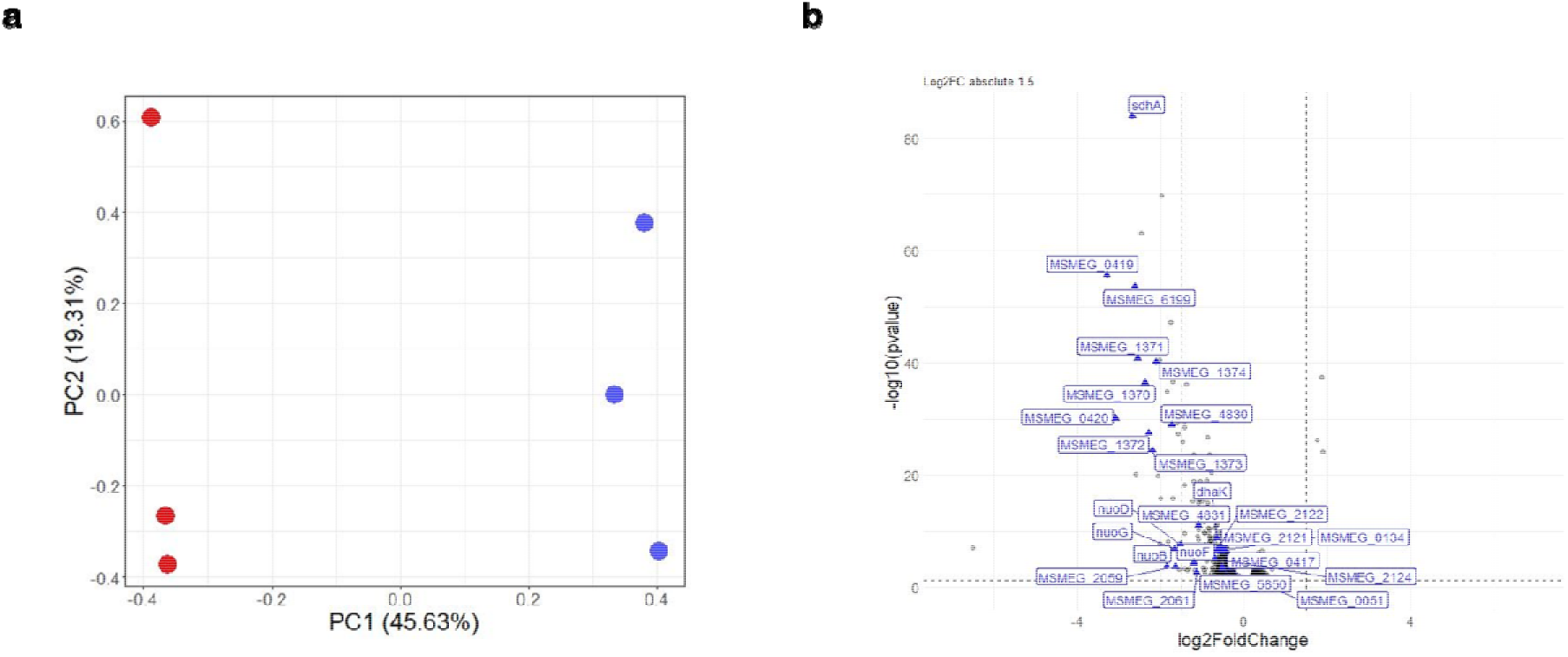
RNA sequencing of empty control vector- and Rv1339-expressing *M. smegmatis* mc^2^155 strains at the mid-log phase of growth. **A.** Principal component analysis of the two bacterial strains. The red colour corresponds to the Rv1339-930 expressing strain, and the blue colour corresponds to the empty control vector-expressing strain. **B.** Volcano plot of significantly differentially expressed genes (p<0.05) in the empty control vector- and Rv1339-expressing *M. smegmatis* mc^2^155 strains. The labelled genes have previously been shown to be regulated by cAMP. The experiments were performed in technical triplicates.

*M. smegmatis* mc^2^155 encodes two Crp genes (Crp 1 and Crp 2^50^): Crp 1 shows greater homology to Crp_MT_, and Crp 2 exhibits higher homology to Cmr. A previous microarray study investigated the effect of Crp 1 deletion or Crp 2 overexpression^51^ on the transcriptome. Attempts to delete Crp 2 were repeatedly unsuccessful, which suggests that this gene is essential for *M. smegmatis* mc^2^155. The deletion of Crp 1 significantly increased the expression of 54 genes, and many of these genes are involved in bioenergetic processes, such as ATP production, succinate dehydrogenases, NADH dehydrogenases and a variety of ABC transporters^51^. This finding suggests that Crp 1 mediates the repression of these genes and exerts some control over bacterial bioenergetics. Furthermore, in the Crp 2-overexpressing strain, 20 genes displayed significantly increased (1.5- to 6-fold) expression^51^. This finding suggests that Crp 2 promotes the expression of these genes, which include several succinate dehydrogenase subunits and the resuscitation-promoting factor rpfE. Another study also suggests that Crp 1 and Crp 2 play different roles in negative and positive gene regulation, respectively^50^.

First, we sought to confirm the previously published data on the Crp regulon in *M. smegmatis* mc^2^155 and to offer further insights into the mechanisms of Crp-mediated gene regulation. Alterations in the levels of genes encoding essential bioenergetic processes could also explain the altered antimicrobial susceptibility because the bacterial bioenergetic state has previously been linked to AMT and AMR^52,53^.

In Rv1339-expresing strains, which exhibited significantly decreased cAMP levels, 387 genes were differentially expressed (p<0.05, the full gene list is shown in Supplementary Table 3). Most of the 387 genes displayed decreased expression (312), and 40 genes were downregulated by at least 1.5-fold (p<0.05) (Figs. 4a and 4b). Conversely, only five genes were upregulated by at least 1.5-fold (p<0.05). Genes that were previously identified in the Crp regulons are indicated with blue labels (Fig. 4b).

Our data on the regulated genes are in agreement with observations from the Crp 1 and Crp 2 microarray study^51^. This included genes involved in most of the bioenergetic processes that were previously identified to be in the Crp regulon, which also confirmed the proposed role of Crp in bioenergetic regulation (Supplementary Table 3). Effectively, 14 of the 40 significantly downregulated genes identified in our study were previously shown to be regulated by Crp 1 and Crp 2. This set of genes included three of four genes encoding the Sdh1 succinate dehydrogenase (MSMEG_0417-0420) operon^54^, which shows decrease in expression by 3- to 4-fold. In *M. tuberculosis*, H37Rv Sdh1 is essential for the survival and maintenance of the proton motive force (PMF) under hypoxic infection conditions^55,56^. Our data also included six of 14 genes encoding the Nuo NADH dehydrogenase operon (all exhibited changes in expression of approximately 2.5-fold) (Fig. 4b). These genes form part of complex II and I of the electron transport chain (ETC). Overall, these data suggest that a decrease in the intracellular cAMP levels alters the bioenergetics of *M. smegmatis* mc^2^155.

### A decrease in the intracellular cAMP levels compromises bacterial bioenergetics

A multipronged approach was used to investigate the effects of decreased intracellular cAMP levels on bacterial bioenergetics. First, the extracellular acidification rates (ECAR) and oxygen consumption rates (OCR)^57–59^ (Fig. 5) of the empty control vector-, Rv1339- and Rv1339 D180A-expressing *M. smegmatis* mc^2^155 strains were determined in the presence and absence of a carbon source. The activities measured by the ECAR include carbon catabolism and the tricarboxylic acid cycle (TCA) due to the production of H^+^ as a result of glycolysis or from NADH/H^+^ synthesis. NADH is a reducing equivalent/electron donor that feeds electrons into the menaquinone pool (MK) and then the ETC. Succinate is also a reducing equivalent/electron donor, but succinate dehydrogenase activity is not measured by the ECAR. The OCR is a readout of the activity of the ETC. To monitor both the ECAR and OCR, Seahorse XFP analysis was performed with the three *M. smegmatis* mc^2^155 strains used in this study. A protocol was designed based on previous studies in the literature^58,59^ (Figs. 5a and 5b). After three cycles of measurements over the course of 20 minutes (1 cycle = 4 minutes of mixing and 2 minutes of reading), the basal OCR and ECAR in the absence of a carbon source were measured to gain an understanding of the basal bioenergetic capacity. The data clearly demonstrate that Rv1339-expressing bacteria exhibited a higher basal OCR and an increased ECAR over the course of the three initial measurements (~20 minutes), which indicates that the bioenergetic machinery was performing at increased levels in the Rv1339-expressing strain, even in the absence of a carbon source. After 20 minutes, glycerol and glucose were injected to a final concentration of 0.2%, which is consistent with the carbon concentration in the bacterial culture media used in all other experiments. Once the carbon sources were injected, all bacterial strains displayed increases of more than 10-fold in their OCR and ECAR. However, in the Rv1339-expressing bacteria, the levels of OCR and ECAR throughout the rest of the experimental period were significantly higher than those of the empty control vector-expressing strain. This finding indicates that even in the presence of carbon sources, the bioenergetic machinery of the Rv1339-expressing bacteria is forced to work much harder than that of the empty control vector-expressing strain. The Rv1339 D180- expressing bacteria, which produces an inactive PDE, displayed similar OCR and ECAR levels as the empty control vector-expressing strain throughout the assay, and the differences between the strains were not statistically significant.

**Figure 5:**
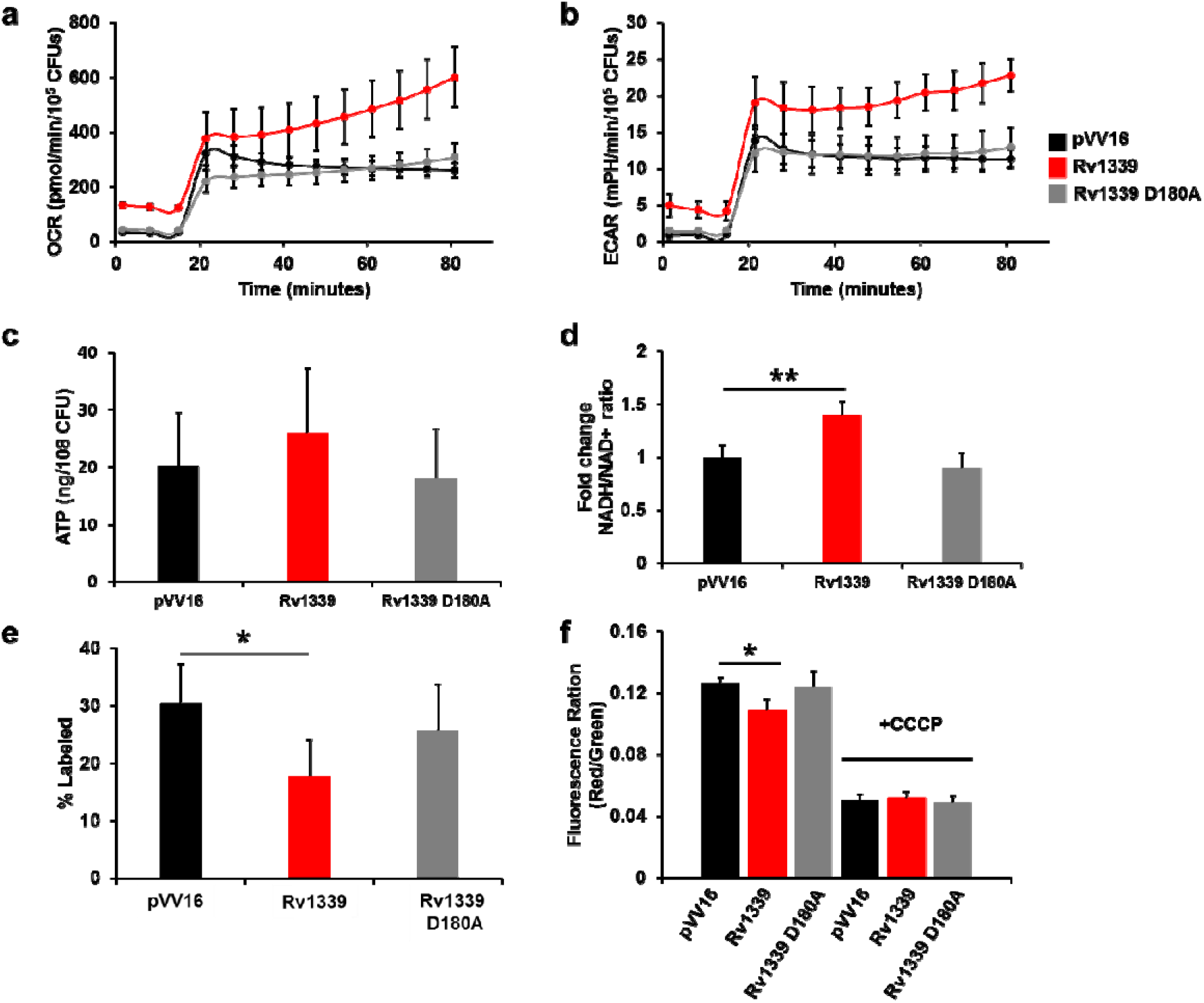
Bioenergetics of empty control vector-, Rv1339- and Rv1339 D180A- expressing *M. smegmatis* mc^2^155 strains. **a**. Oxygen consumption rate of the empty control vector- and Rv1339-expressing *M. smegmatis* mc^2^155 strains. **b**. Extracellular acidification rate and oxygen consumption rate of empty control vector- and Rv1339-expressing *M. smegmatis* mc^2^155 strains. **c**. ATP levels measured at the mid-log phase of bacterial growth. d. Fold change in the NADH/NAD+ ratio measured at the mid-log phase of bacterial growth. Succinate turnover measured by ^13^C-label incorporation. **f**. Membrane potential measurements at the mid-log phase of bacterial growth. The statistical analysis was performed with a two-tailed Student’s t-test. * p<0.05 and **p<0.001. The data are presented as the means±SDs from three biological replicates and three replicates.

Increased OCR and ECAR activity should correlate with increased production of ATP (from carbon catabolism and/or the TCA cycle and/or ETC activities), which is the precursor of cAMP and a critically important energy source for a multitude of bacterial activities. Similarly, an increased ECAR should correlate with increased NADH levels because NADH is produced by carbon catabolism and TCA cycle activity^58^. To confirm this hypothesis, we quantified the intracellular ATP levels and the fold changes in the NADH/NAD+ ratio in the empty control vector-, Rv1339- and Rv1339 D180A-expressing *M. smegmatis* mc^2^155 strains (Figs. 5c and 5d). Surprisingly, despite significant differences in the bioenergetics rates, no differences in the ATP levels were found among the three strains (Fig. 5c). Although Aung *et al* found that Crp 1 regulates the alternative mycobacterial cytochrome *bd* and ATP synthase components in their microarray study^51^, no increases in these genes were found in our dataset (Supplementary Table 3). The NADH-to-NAD+ ratio was then determined for the three strains (Fig. 5d). The NADH/NAD+ ratio serves as an indicator of carbon catabolism and/or TCA cycle activity. NADH, which is produced from NAD+, feeds electrons into the ETC and maintains bacterial bioenergetics^52,58,60–62^. The NADH/NAD+ ratio was increased by 0.5-fold in the Rv1339-expressing strain compared with the empty vector control (p<0.01), which indicates an increase in carbon catabolism.

To investigate the changes in carbon catabolism activities, the turnover of serine, which can be used as a readout of glycolytic activity^63^, was measured by [U-^13^C_6_]-glucose and [U-^13^C_3_]- glycerol stable isotope labelling (Supplementary Fig. 2c). This analysis clearly showed that Rv1339-expressing bacteria exhibited increased ^13^C incorporation (e.g., m+3 of serine is increased by 10% in the Rv1339- compared with the empty control vector-expressing strain) (Supplementary Fig. 2c and Supplementary Table 5). This finding indicates an increased turnover of serine and thus increased glycolytic activity. Additionally, the *sdaA* gene exhibited a 0.35-fold decrease in expression. This gene encodes *L*-serine ammonia lyase, which is an enzyme that can convert serine to pyruvate. Concurrently, the decreased *sdaA* expression might indicate compensation for the reduced enzyme activity of this pathway through an increase in substrate production. This finding suggests that this enzyme might replenish the pyruvate pool, and this alteration could produce more NADH and ATP and thus contribute to the increased bioenergetic activity.

As mentioned earlier, a decrease in intracellular cAMP leads to a decrease in *sdh1* expression, which potentially results in a decrease in the succinate turnover. To test this hypothesis, we conducted ^13^C isotopologue analysis (Fig. 5e). Sdh1 catalyses the conversion of succinate to fumarate^54^; therefore, decreased Sdh1 activity would be indicated by decreased turnover of succinate. The rate of succinate turnover was indeed decreased by 33% in the Rv1339-expressing bacteria compared with empty control vector-harbouring bacteria. This decrease was not observed in the Rv1339 D180-expressing strain, which produces a catalytically inactive enzyme (Fig. 5e and Supplementary Fig. 2b), and these findings thus confirmed that the effect observed was due to the cAMP PDE activity of Rv1339.

The above-described findings further reinforce the increased bioenergetics observed in the Seahorse data because an increased ECAR might be indicative of increased carbon catabolism or increased TCA cycle activities. Due to the increased OCR, it is unlikely that the increase in the NADH/NAD+ ratio is due to a reduced influx of electrons from NAD+ into the ETC. Moreover, the membrane potential of the pVV16-, Rv1339- and Rv1339 D180A-expressing *M. smegmatis* mc^2^155 strains was measured. The decrease in intracellular cAMP levels in the Rv1339-expressing strain was accompanied by a 15% decrease in the membrane potential compared with the levels found in the empty control vector-expressing strain (Fig. 5f), and this decrease was not observed in the Rv1339 D180-expressing bacteria (Fig. 5f). The membrane potential is a key determinant of the PMF, and it is known that the PMF drives the proton gradient across the bacterial membrane required for ATP synthesis^64,65^. A decrease in the membrane potential would impact the PMF, and in combination with the observed 33% decrease in succinate turnover, the observed decrease in the membrane potential indicates that bacterial bioenergetics is compromised, which likely has an impact on the membrane permeability.

### Reduced intracellular cAMP levels lead to increased bacterial permeability

The production, maintenance and alteration of the cell wall composition is an ATP- and resource-intensive task performed by bacteria^36^, and these processes are also crucially important in modulating the susceptibility to antimicrobials by maintaining bacterial bioenergetics during infection^53^. To investigate the effect of decreased cAMP levels and compromised bacterial bioenergetics on the integrity of the cell wall, two independent approaches were used: crystal violet and Hoechst dye 33342 uptake assays^66^ (Supplementary Fig. 3). Crystal violet is a toxic dye, and bacteria with higher permeability show greater uptake of this dye. The Rv1339-expressing bacteria showed decreased growth on crystal violet-containing agar compared with the empty control vector-expressing strain (Supplementary Fig. 3a), which indicated an increase in permeability. This finding was confirmed by measuring the uptake of a DNA binding dye, Hoechst 33342, by bacteria grown in liquid culture. The Rv1339-expressing bacteria showed increased incorporation of the dye, which indicated increased permeability (Supplementary Fig. 3b).

### Untargeted metabolomics reveals that reductions in the intracellular cAMP levels increase the plasticity of peptidoglycan synthesis

As mentioned previously, a decrease in the intracellular cAMP levels alters cellular bioenergetics, central carbon metabolism and permeability. To elucidate the mechanism underlying this increased permeability, an untargeted metabolomics analysis was performed (Fig. 6), and this analysis (Fig. 6a) clearly showed that the decreased cAMP levels in the Rv1339-expressing strain resulted in substantial alterations in the levels of compounds that were later confirmed to be meso-2,6-diaminoheptanedioate (m-DAP – 2.2-fold), lysine (1.2-fold), *D*-alanyl-*D*-alanine (1.5-fold), *N*-acetyl lysine (1-fold), *N*-acetyl-alpha-glucosamine-6-phosphate (4-fold) and tetrahydrodipicolinate (4-fold) (Fig. 6b). These alterations were not observed in the Rv1339 D180A-expressing bacteria (Fig. 6b). Interestingly, all of these metabolites are involved in peptidoglycan biosynthesis, which is an ATP-intensive process^36^. The observed alterations in the metabolite pool sizes were validated by measuring their turnover through the incorporation of ^13^C at half-doubling time. We found that the turnover of *m*-DAP, *D*-ala-*D*-ala, *N*-acetyl-lysine and lysine was drastically altered (Supplementary Fig. 4), and this altered turnover of these metabolites and the changes in the metabolite pool sizes indicates greater plasticity in the peptidoglycan layer. Both lysine and *m*-DAP displayed decreased metabolite pool sizes, despite increased turnover (Fig. 4b and Supplementary Fig. 4). *m*-DAP is a key component of peptidoglycan, and the fact that the absolute levels of *m*- DAP were decreased despite an increase in turnover suggests that more *m*-DAP is being produced and then consumed. The finding that the lysine metabolite pool size was also decreased despite an increased turnover might suggest that although more *m*-DAP is being produced, the muralytic enzyme LysA^67^ might break down more *m*-DAP from peptidoglycan into lysine, thereby indicating increased production and degradation of peptidoglycan and therefore increased plasticity. DAP crosslinks provide stability to the cell wall, and thus, increased plasticity likely indicates increased instability^67^. The metabolite pool size of tetrahydrodipicolinate, which is the precursor of *m*-DAP, displayed a 4-fold increase, which might compensate for the increased turnover of *m*-DAP. The production of *m*-DAP has been shown to be essential in mycobacteria because its disruption leads to decreased stability of peptidoglycan^67,68^. Furthermore, *D*-alanyl-*D*-alanine and *N*-acetyl-alpha-glucosamine (GlcNAc) are also key components of peptidoglycan, and GlcNAc represents half of the glycan strand. The pool sizes of both of these metabolites were 1.5-fold and 4-fold higher, respectively. Additionally, the turnover of *D*-ala-*D*-ala was decreased, which suggested that the increase in the metabolite pool might not be the result of increased synthesis but perhaps degradation. Taken together, these findings further support the hypothesis that decreased intracellular cAMP levels result in increased production and degradation of peptidoglycan. An increased plasticity of peptidoglycan and decreased stability would likely lead to negative effects on bacterial permeability to antimicrobials.

**Figure 6:**
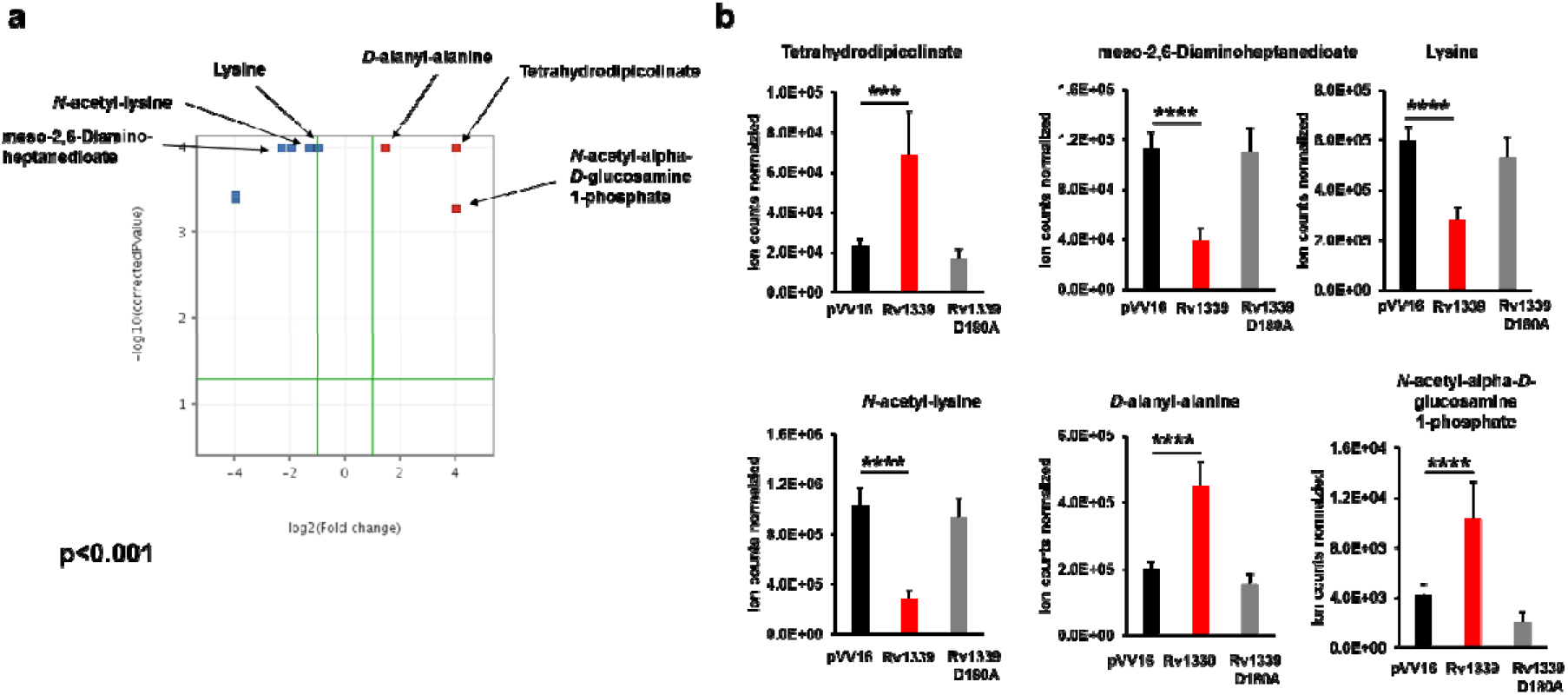
Untargeted metabolomics analysis reveals that peptidoglycan synthesis is altered in *M. smegmatis* mc^2^155 pVV16::rv1339. **a**. Volcano plot displaying the differential abundance of metabolites in the empty control vector- and Rv1339-expressing *M. smegmatis* mc^2^155 strains. **b**. Abundances of metabolites in the three strains investigated in this study. The experiments were performed in biological duplicate and technical quadruplicate. Unpaired two-tailed Student’s t-tests were used to compare the data. *p<0.05, **p<0.01, ***p<0.001 and ****p<0.0001. The data are presented as the means±SDs from two biological replicates and three replicates.

## DISCUSSION

Although the involvement of cAMP in antimicrobial susceptibility in pathogenic gram-negative bacteria has been previously elucidated^6,31^, our data provide the first demonstration of the consequences of reducing the cAMP pool in mycobacteria without deleting any cAMP synthesis genes. Decreasing the cAMP levels using this approach might be a useful tool for investigating this signalling system that has less background noise than previous approaches used in the field^17,69^.

The expression of Rv1339 decreased the intracellular cAMP levels and increased the bacterial susceptibility to antimicrobials with widely different modes of action. Along with decreased cAMP levels, we observed decreased expression of key components of mycobacterial bioenergetics and increased permeability due to increased peptidoglycan plasticity.

Indeed, cAMP signalling has previously been shown to regulate the susceptibility of *Salmonella typhimurium* and uropathogenic *E. coli* to antimicrobials^6,7,31^. Studies have shown that this result in *Salmonella typhimurium* is due to increased permeability for antimicrobials^6,7^. The aforementioned bacteria are both gram-negative, and our data provide first elucidation of the link between cAMP and permeability in an organism with a high prevalence of peptidoglycan, which forms a thick portion of the complex mycobacterial cell wall^36^. This layer around the bacteria at least partly explains why mycobacteria are innately tolerant to antimicrobials^36^, and thus, it might be unsurprising that the increased peptidoglycan plasticity observed in this study could underpin the increased susceptibility of the bacteria. Previous studies have shown that the utilization of trehalose, which is the sugar portion of several mycobacterial cell wall lipids, underpins the development of transient antimicrobial tolerance and permanent resistance^53^. This finding further supports the involvement of alterations in the mycobacterial cell wall composition in the modulation of antimicrobial susceptibility observed in this study.

Previous studies published in the literature have linked the ATP levels to the development of persistence and tolerance to antimicrobials. In *E. coli*, persister formation and tolerance are thought to be dependent on the ATP levels^70^. Lower levels lead to a slow-down of bacterial processes under antibiotic treatment and thus increased tolerance to the antimicrobial (a fluoroquinolone). Similarly, in *Staphylococcus aureus*, tolerance and persister formation during exponential growth are correlated with reduced stationary-phase-like ATP levels^71^. A more recent study that investigated *Salmonella enterica* supports the hypothesis that slow growth mediates tolerance and persistence but refutes the presumed requirement for low ATP levels^72^. Instead, slow growth and tolerance can be induced even with high ATP levels, and slow growth drives this transition^72^. The uncoupling of tolerance from decreased ATP levels is supported by our data, which do not show any significant alterations in the ATP levels but increased bacterial susceptibility to antimicrobials.

Even though Rv1339-expressing bacteria exhibited a higher OCR and an increased ECAR, we did not observe any increase in the ATP levels in the mid-log phase. Although this finding was unexpected, other studies have shown that even without significant changes in the absolute ATP levels, bacterial metabolism and bioenergetics can still be significantly altered^73^. Furthermore, we found decreases in the expression of 33 genes encoding ATP-binding proteins, including numerous ABC transporters, a 30% decrease in growth and increased plasticity in peptidoglycan synthesis, which likely indicates reduced stability. The proteins encoded by these genes are ATP binders, and growth and peptidoglycan biosynthesis are ATP-dependent processes^36,74,75^. In summary, it appears that forcibly decreasing the cAMP levels is sufficient to place stress on the ATP pool. We hypothesize that to maintain the essential membrane potential and bioenergetic processes, e.g., the succinate dehydrogenase activity of Sdh1, bacteria are forced to commit a deleterious quantity of resources towards maintenance of the cAMP levels.

There is a precedence for this hypothesis: during macrophage infection, *M. tuberculosis* and *M. bovis* BCG bacteria increase their cAMP levels by 50-fold^76^. Because the cAMP levels in *M. tuberculosis* are innately high, it is likely that a sudden increase in the cAMP levels would put strain on the ATP pool.

Interestingly, the literature describes additional circumstantial evidence that supports this hypothesis. Single point mutations in Rv1339 have been found to confer resistance to the entire class of imidazolidines (IMPs), which are ATP-depleting compounds that are structurally related to Q203^77^. The characterization of the IMPs suggests that these proteins target QcrB, a proton-pumping component of the terminal cytochrome oxidase (complex III of the ETC). *M. tuberculosis* treated with IMPs exhibit dose-dependent decreases in the ATP levels and disruption of the intrabacterial pH. Mutations in QcrB also lead to resistance^77^. All single point mutations reported in Rv1339 cluster around the metal-binding site or are predicted to change the structure of the enzyme. If these mutations abrogate the activity of Rv1339 and decrease the demand on the ATP pool, these effects might compensate for the effects of ATP depletion.

Although *M. tuberculosis* produces cAMP to intoxicate host macrophages and prevent immune-mediated killing^78,79^, a 50-fold increase in the bacterial cAMP level might represent adaptation to the host macrophage environment. This hypothesis is supported by the presence of 16 cAMP-producing enzymes that also contain sensory domains in *M. tuberculosis* H37Rv^8,14^. Based on our data, which showed that decreases in the cAMP levels by just 3-fold resulted in increased permeability, it is tempting to speculate that increased cAMP levels might play a role in the adaptation of the bacterial cell wall composition to reduce permeability and allow improved survival in the harsh environment of the host macrophage. Several studies have linked cAMP with mediating bacterial adaptation in response to the NaCl levels found within the host macrophage, which only further supports the role of cAMP in the essential adaptation to the host environment during infection^30,80^.

Further evidence supporting the importance of cAMP signalling in adaptation to and survival in the host environment has been found during *M. tuberculosis* infection. Host-derived cholesterol is the preferred carbon source of *M. tuberculosis* during host infection^81,82^. Successful infection requires the ability to metabolize cholesterol and detoxify downstream products^83^. Detoxification of the cholesterol catabolism product propionyl-CoA requires the expression of genes that are known to be regulated by cAMP^84,85^. Additionally, a screen of mycobacterial cholesterol-targeting small molecules identified several promising compounds with significant bactericidal efficacy against intracellular *M. tuberculosis* during macrophage infection. Although the mode of action has not been fully characterized, treatment with the compounds led to drastically increased cAMP levels via activation of the bacterial adenylate cyclase Rv1625c^86,87^. Point mutations in Rv1625c abrogated this increase and conferred resistance to the compounds. The fact that cAMP appears important to both cholesterol metabolism and downstream detoxification makes cAMP signalling a promising target because bacteria would find it more difficult to evolve resistance under these conditions.

Overall, our data suggest that targeting cAMP signalling could lead to impairments of bacterial bioenergetics and viability during infection. We also provide circumstantial evidence showing that decreasing the cAMP levels using a conserved actinobacteria phosphodiesterase enzyme can prevent the development of resistance to streptomycin and drastically increase the efficacy of ethambutol. These effects are likely a result of increased permeability, which can be explained by compromised peptidoglycan integrity. Therefore, there is a strong incentive to further investigate the validity of targeting the cAMP signalling system in mycobacteria, and this further study might provide insights for increasing the susceptibility of these bacteria to antimicrobial treatment or inhibiting the development of resistance.

## MATERIAL AND METHODS

### Bacterial strains and culture conditions

*M. smegmatis* mc^2^155, parental *M. bovis* BCG and *M. bovis* BCG ∆*rv0805* were used in this study. The mycobacteria were cultured to the mid-exponential phase in 7H9 liquid medium supplemented with 0.5 g/l Fraction V (bovine serum albumin), 0.05% tyloxapol, 0.2% dextrose, 0.2% glycerol, and 10 mM NaCl. For the time-to-kill assays, crystal violet assays and metabolomic profiling studies, the mycobacteria were cultured on 7H10 agar supplemented with 0.5 g/l Fraction V (bovine serum albumin), 0.2% dextrose, 0.2% glycerol and 10 mM NaCl. Throughout the study, the mycobacteria were cultured in static or shaking incubators at 180 rpm and 37°C. The *M. bovis* BCG strains were kindly provided by Prof Sandhya Visweswariah from the Indian Institute of Science, Bangalore, India.

### Plasmid constructs

*rv1339* was cloned using primers that allow isothermal assembly (Gibson) using the pVV16 vector containing kanamycin and hygromycin resistance cassettes.

The following primers were used:

open_pVV16_forward
5’-AAGCTTCACCACCACCACCACCACTGACAG-3’
open_pVV16_reverse
5’-CAT ATG GAA GTG ATT CCT CCG GAT CGG GGA TG-3’
Rv1339_pVV16_forward
5’-CCGATCCGGAGGAATCACTTCCATATGCGTCGATGTATTCCGCATCGTT-3’
Rv1339_pVV16_reverse
5’-GTCAGTGGTGGTGGTGGTGGTGAAGCTTGCCGGCTCGCCGGACTTCG-3’
Rv1339_D180A_forward
5’-CTCGCGGCGTCGCCGTTTTCCTCTGC-3’
Rv1339_D180A_reverse
GCAGAGGAAAACGGCGACGCCGCGAG

All insert and vector PCR products were cleaned with Qiagen PCR clean-up DNA spin columns, and for site-directed mutagenesis, the template DNA was degraded by restriction digestion with DpnI (NEB). The fragments were joined at 50°C for 15 minutes using HiFi Builder Assembly Master Mix. DH5-α chemically competent cells were used as the plasmid library strain, plated on LB agar containing kanamycin at a final concentration of 60 μg ml^−1^ and incubated overnight at 37°C. Colonies were screened using KAPA2G Robust HotStart ReadyMix and a colony PCR protocol. The positive colonies were mini-prepped, and the subsequent construct was used to transform *M. smegmatis* mc^2^155 bacteria by electroporation. The transformants were plated on 7H10 agar using hygromycin (50 μg ml^−1^) and kanamycin (25 μg ml^−1^) as the selection markers.

### Preparation of clear soluble lysate

Cultured *M. bovis* BCG cells were aliquoted into 50-ml Falcon tubes under a class II hood and sealed. The cells were then centrifuged at 3,000 × g. All centrifugations were performed at 4°C for 5 minutes. The supernatant was discarded, and the pellet was washed three times with 20 mM Tris-HCl (pH 7.5), 50mM NaCl containing protease inhibitor (Sigma fast). One millilitre of washed bacteria was aliquoted into 2-ml microtubes containing 200 μl of 0.1-mm acid-washed soluble lysate, and 200 μl of the resulting mixture was aliquoted into Eppendorf tubes. These aliquots were then frozen in liquid nitrogen and maintained at −80°C for further analysis.

### Chromatography conditions for cAMP PDE activity enrichment

All column purifications of *M. bovis* BCG lysates were performed using ÄKTA systems at 4°C. The systems were operated using Unicorn manager software. Capto Q is an anion-exchange column that separates proteins across a NaCl gradient. Low (30 mM) and high (600 mM) concentrations of NaCl with 20 mM Tris-HCl (pH 7.5) were used as buffer A, and size-exclusion chromatography was performed using a 24-ml Superdex 200 10/300 column (GE Healthcare) with 20 mM Tris (pH 7.5) and 30 mM NaCl.

### PDE activity assay

Ten microlitres of cell lysate/fractionated lysate/purified protein was incubated at 37°C for 16 hours with 2 μl of water, 2 μl of 10x PDE buffer (20 mM MgCl_2_, 20 mM Tris-HCl pH 9.0 and 100 mM NaCl) and 6 μl of 100 mM (for TLC) or 25 mM (for LC-MS) cyclic AMP at 10 mM or water (as a control). For the 5’-AMP controls, 6 μl of 5’-AMP (10 mM) was added at the same concentrations of cAMP. All PDE reactions were performed in Eppendorf tubes.

### Thin-layer chromatography

Silica gel 60 F254 TLC plates were cut to the required size of less than 15 × 10 cm. The 50-ml mobile phase consisted of 70:30 ethanol/H_2_O 0.2 M ammonium bicarbonate. After migration, the plates were revealed by shearing a solution of 5% phosphomolybdic acid dissolved in ethanol and heating at 150°C for 10 minutes.

### Identification of the PDE candidate by proteomics analysis

The fraction displaying PDE activity, as determined by TLC after Capto Q and SEC column purifications, and a fraction on either side were run on an SDS gel as described previously and stained with InstantBlue™ Coomassie stain. The gel lanes were cut, reduced, alkylated and digested with trypsin using the ProteaseMAX Surfactant protocol. The samples were then loaded and run on a SYNAPT Q-ToF mass spectrometer. The observed proteins were correlated to the protein IDs and potential annotations in the UniProt database.

### Bioinformatics analysis

The sequences were aligned with MUSCLE^88^. The best-fit amino acid substitution model for the alignment was LG+R6, which was found using ModelFinder^89^. The trees were established using IQ-Tree (1.6.11^90^) with 100 bootstrap replicates and visualized using Dendroscope^91^. Selected solved 3D structures are shown with a diamond and labelled with their PDB accession code. The characterized members of the families are the following (their UniProt and PDB accession codes are shown in brackets): *Aliivibrio fischeri* cpdP (Q56686), *Yersinia pestis* cpdP (Q8ZD92), *Saccharomyces cerevisiae* PDE1 (P22434; 4OJV), *Escherichia coli* phosphotriesterase homology protein (PhP; P45548; 1BF6), *Homo sapiens* zinc phosphodiesterase ELAC protein 1 (Q9H777; 3ZWF), *Escherichia coli* ribonuclease BN (Rbn; P0A8V0; 2CBN), *Mycobacterium tuberculosis* ribonuclease Z (P9WGZ5), *Bacillus subtilis* ribonuclease Z (P54548; 1Y44), *Bacillus subtilis* metallo-hydrolase YhfI (O07607; 6KNS), *Streptomyces coelicolor* (SCO2908), *Corynebacterium glutamicum* CpdA a.k.a. Cgl2508 (Q8NMQ7), and *Mycobacterium tuberculosis* H37Rv Rv1339 (P9WGC1).

### Bacterial growth assay

Ten-millilitre square bottles were prepared by adding 10 ml of 7H9 medium and incubating in a static incubator to allow the media to reach a temperature of 37°C. A preculture was prepared by adding 100 μl of bacteria stored at −80°C to 10 ml of culture medium. Once the preculture reached the mid-log phase, the growth curve was initiated. Specifically, 125-ml conical flasks containing 25 ml of 7H9 medium were inoculated with mycobacteria (from the preculture) at an OD_600_ of 0.001 and then incubated at 37°C in a shaking incubator set to 180 rpm. The OD_600_ was monitored over 80 hours.

### His-tagged protein analysis of mycobacteria

To confirm the expression of Rv1339 or Rv1339 D180A, all strains of *M. smegmatis* were grown in Middlebrook 7H9 broth, as previously described. The bacterial cultures were centrifuged at 3,000 × g and 4°C for 10 minutes to harvest the bacteria. The pellets were washed three times with 20 mM Tris-HCl and 50 mM NaCl (pH 7.5) and then centrifuged at 3,000 × g and 4°C for 10 minutes. The pellets were suspended in 1.5 ml of lysis buffer containing protease inhibitor cocktail (Sigma-Aldrich, USA) and transferred into an O-ring tube containing 100 μl of 0.1-mm acid-washed zirconia beads. All the samples were ribolysed three times with an MP FastPrep®-24 homogenizer (MP pharmaceuticals) for 60 seconds. The samples were then centrifuged at 11,000 × g for 10 minutes. The supernatant was transferred to an Eppendorf tube, and the protein concentration was determined using a Nanodrop Lite Spectrophotometer (Thermo Fisher Scientific) and BCA assay. All the samples were diluted with lysis buffer to ensure equal protein concentrations, added to 5X SDS loading buffer (0.25 M Tris-HCl pH 6.8, 15% SDS, 50% glycerol, 25% β- mercaptoethanol, and 0.01% bromophenol blue) and boiled at 100°C for 5 minutes. Subsequently, 50 μg of each sample was loaded onto the SDS-PAGE gel, and the gel was then run at 150 V and 40 mA for 1 hour in a tank filled with SDS running buffer (3% Tris, 14.4% glycine, and 1% SDS).

After running the SDS-PAGE, the gel was transferred onto a nitrocellulose membrane (Novex, USA) by running at 20 V and 400 mA for 1 hour at room temperature. The tank was filled with a transfer buffer containing 10X SDS (3% Tris, 14.4% glycine, and 1% SDS), methanol, and ddH_2_O at a ratio of 1:2:7. The non-specific sites on the nitrocellulose membrane were blocked overnight with 10 ml of 5% skimmed milk (Marvell, UK) suspended in buffer composed of 0.06% Tris base, 0.88% NaCl, and 0.1% Tween 20 (pH 7.5). The membrane was washed with the buffer, incubated for 1 hour with mouse HRP α-His Tag antibody (1:1000 in 5% milk) (BioLegend, USA) and developed. The membrane was then washed using the same above-mentioned steps, stripped with ReBlot Mild Plus (Sigma) and incubated with an anti-mtbGroEL2 antibody (BEI resources). The membranes were washed as previously described and incubated with a goat anti-mouse antibody conjugated to horseradish peroxidase (1:5000 in 5% milk), which served as the loading control. Rv1339 protein was visualized with a FujiFilm LAS-3000 Image Reader and the Amersham™ ECL™ Western Blotting Analysis system (GE Healthcare, UK).

### Determination of the minimal inhibitory concentration by a resazurin microtiter assay (REMA)

The minimal inhibitory concentration (MIC_90_) was determined using the REMA according to the NCCLS guidelines^92^ and Palomino *et al*. (2002)^93^. Antibiotic stock solutions of the tested compounds were prepared to yield the target concentrations for testing, as shown in Table 1. The microdilution assays were performed in 96-well plates. Two-fold serial dilutions were used to obtain the final drug concentration, which ranged from 0.016 to 4 μg/ml (ethambutol) and 0.008 to 2 μg/ml (streptomycin). To obtain the bacterial inoculum, the empty vector- and Rv1339-expressing *M. smegmatis* strains were grown to the mid-log phase (OD_600_ ~ 0.5) and diluted 1:1000 with 7H9 broth. Fifty microlitres of the standardized bacterial inoculum and 50 μl of 7H9 broth were added to each well of a 96-well plate. The plates were incubated for 48 hours at 37°C with 5% CO_2_. Subsequently, 30 μl of 0.01% resazurin was added, and the mixture was maintained at 37°C with 5% CO_2_. After colour development for 24 hours, the wells were read. The MIC_90_ was defined as the lowest concentration that inhibited the growth of 90% of bacteria.

### Time-to-kill assays

The empty vector- and Rv1339-expressing *M. smegmatis* strains were grown to the mid-log phase (OD_600_ ~ 0.5) and diluted 1:100 to yield ~10^5^-10^6^ CFUs/ml in Middlebrook 7H9 medium. Antibiotics were added to each sample at defined concentrations, and the bacterial samples were collected before addition of the antibiotic and 6, 24, 30, 48 and 72 hours after antibiotic challenge. To determine the number of viable cells, the CFUs were determined through serial 10-fold dilutions using 20 μl of culture and 180 μl of 7H9 broth. Twenty microlitres of each dilution was plated onto Middlebrook 7H10 agar (Sigma-Aldrich, Germany). All the plates were incubated at 37°C for 4 days before the colonies were counted.

### Permeability assay using Hoechst staining

The mycobacterial strains were grown in complete 7H9 medium until the exponential phase and diluted to a final OD_600_ of approximately 0.8 with the same medium in a total volume of 5 ml. The bacterial suspension was centrifuged for 10 minutes at 4,000 × g and 4°C, and the supernatant was removed. The pellet was washed twice with sterile cold PBS by suspending the pellet and centrifuging the suspension at 4,000 × g and 4°C for 25 minutes, and the supernatant was removed. The pellet was suspended in 5 ml of complete 7H9 medium. White-bottomed 96-well plates were prepared by adding 200 μl of 7H9 medium to the perimeter to minimize medium evaporation, and 180 μl of the bacterial suspension was then loaded into separate columns of the plate. Blank columns were prepared by adding the same volume of the corresponding media. Hoechst 33342 stain solution was prepared to a final concentration of 25 μM with sterile deionised H_2_O. Twenty microlitres of the stain solution was loaded into each well of the plate, and the loading order was switched between two groups to avoid loading bias. The changes in the fluorescence of each well were monitored using a microwell plate reader (Hidex Sense) during incubation at 37°C with shaking at 600 rpm, and the fluorescence was measured with excitation and emission filters of 355 nm and 460 nm, respectively, every 10 minutes for 180 minutes, which represents a doubling cycle of *M. smegmatis* mc^2^155. The fluorescence data were exported into Microsoft Excel to quantify the changes in the fluorescence levels, and the data from the empty control vector- and Rv1339-expressing *M. smegmatis* mc^2^155 strains were compared.

### Crystal violet spot assay

To evaluate the cell membrane permeability of the bacteria, the spotted assay was performed according to the literature^18,94,95^. The bacteria were grown in Middlebrook 7H9 broth, as described above, to the mid-log phase to obtain an OD_600_ value of 0.5, and serial dilutions were performed (undiluted to 10^−7^). Five microlitres of each dilution was spotted on Middlebrook 7H10 agar (Sigma-Aldrich, Germany) containing 10 μg/ml crystal violet. The agar plates were incubated at 37 °C for 3 days and then photographed.

### Analysis of the membrane potential

*M. smegmatis* strains were grown to an OD of 0.6, centrifuged and suspended in 7H9 media containing 10 mM NaCl. One-millilitre aliquots were collected at various time points. The samples were washed with 7H9 media without albumin, suspended in 1 ml of 7H9 media without albumin and exposed to 15 μM DiOC2 for 20 minutes at room temperature. The samples were then washed with 7H9 media without albumin and suspended in the same media. The fluorescence was monitored using a SpectraMax Gemini XPS plate reader (Molecular Devices): green 488/530 nm and red 488/610 nm. The membrane potential was calculated as the ratio of red to green fluorescence.

### Intracellular ATP measurement

The intracellular ATP content was measured as described by Koul *et al*.^96^ and Parrish *et al*.^97^. The bacterial cells were harvested by collecting 1.5 ml of the bacterial culture and centrifuging at 17,000 × g for 15 minutes. The pellets were maintained at −80°C until analysis. Each bacterial pellet was mixed with 1.5 ml of boiling Tris-EDTA (100 mM Tris-HCl and 4 mM EDTA pH 7.75), and 200 μl of 0.1-mm acid-washed zirconia beads was added. All the samples were lysed twice using the MP FastPrep®-24 homogenizer (MP pharmaceuticals) at 6 m/second for 60 seconds. The samples were then heated at 100°C for 5 minutes and immediately cooled on ice for 10 minutes. The cell debris was removed by microcentrifugation at 13,000 × g and room temperature for 15 minutes, and the supernatants were transferred to new sterile tubes for the ATP assay. The ATP content was determined using the ATP Bioluminescence Assay Kit CLS II (Roche). Subsequently, 50 μl of luciferase reagent was added to 50 μl of the supernatant sample. The reactions were measured in a microwell plate-reading luminometer (Hidex Sense). To obtain the ATP amount per CFU, the bacterial culture was serially diluted and plated on Middlebrook 7H10 agar. The agar plates were incubated at 37°C, and the colonies were counted 4 days after plating.

### RNA extraction and RNA sequencing

*M. smegmatis* mc^2^155 strains were grown in 7H9 liquid medium at 37°C as described above. The bacteria were grown to the mid-log phase (light transmittance at 600 nm of 0.6) and washed twice with cold PBS at 4°C. Total RNA was extracted according to the FastRNA Pro Blue kit manual (MP). For the removal of genomic DNA, the samples were treated once with RNase-free DNase (Promega) for 3 hours at 37°C and purified using RNAeasy columns (Qiagen) according to the manufacturer’s instructions. The RNA quantity and quality were assessed using a NanoDrop™ 1000 spectrophotometer (Thermo Fisher Scientific, Inc.) and an Agilent 2100 Bioanalyzer with an RNA 6000 Nano LabChip kit (Agilent Technologies, Ltd.). All the samples displayed a 260/280 ratio greater than 2.0 and RNA integrity numbers (RIN) greater than 9.1. RNA was sequenced using an Illumina HiSeq4000 sequencer by the Imperial BRC Genomics Facility at Imperial College London. Samples were sequenced to obtain paired-end reads of a minimum of 75 bp in length.

### RNA-seq data analysis

The sequence quality was checked using FastQC (v0.11.8, http://www.bioinformatics.babraham.ac.uk/projects/fastqc/). All the sequences passed quality control, and the paired-end sequences were aligned to the *M. smegmatis* mc^2^155 reference genome (GCF_000015005.1_ASM1500v1 assembly, NCBI; using the Burrows-Wheeler Transform (BWA) sequence aligner (v0.7.17-r1188^98^). The read counts were calculated at the gene level using feature counts (v1.6.0^99^) and the *M. smegmatis* mc^2^155 annotation (GCF_000015005.1_ASM1500v1 assembly, NCBI). The sequences were also aligned to the *M. tuberculosis* H37Rv reference genome (GCF_000195955.2_ASM19595v2 assembly, NCBI) and annotated using the *M. tuberculosis* H37Rv annotation (GCF_000195955.2_ASM19595v2 assembly, NCBI) to determine the levels of *rv1339* in all the samples. Genes with low counts in all the samples were removed before the sample quality was evaluated through principal component analysis (PCA) using normalised counts and the prcomp function in base R (v3.6.2). Differential gene expression analysis was performed using the DESeq2 Bioconductor package (v1.26.0^100^), and in this analysis, the Rv1339-expressing strain was compared with the empty control vector-expressing strain. Genes were corrected for multiple testing using the Benjamini-Hochberg (BH) method, and significant genes were identified using a false discovery rate (FDR) cut-off of < 0.05 and an absolute log2-fold change of 1.5.

### Bioenergetics analysis using Seahorse XFP

The standard Middlebrook 7H9 recipe was prepared according to the literature^58,59,101^. The carbon source supplement (glycerol and glucose) was prepared to a final concentration of 2% in ddH_2_O and filtered to ensure that it was sterile.

A Seahorse XFP analysis was performed using methods similar to those performed previously. All reagents were purchased from Agilent Technologies, and all work was performed in a laminar-flow hood. The day before the assay was to be run, the Seahorse XFP cartridge probes were hydrated by filling the utility plate and all border wells with 200 μl of sterile ddH_2_O per well. This utility plate-cartridge unit was then incubated in an airtight container overnight.

A minimum of 2 hours before the assay was run, the ddH_2_O was removed and replaced with XFP calibrant solution, and the cartridge was returned to the 37°C incubator. Subsequently, 20 μl of each substance to be injected was placed in the relevant injection port, and during this process, care was taken to ensure that none of the sample was stuck to the sides of the injection port. In the assays performed in this study, the only substance injected was glucose-glycerol, which was injected to a final concentration of 0.2%. Separately, each well of the bacterial cell plate was coated with 90 μl of sterile poly-D-lysine (PDL) and allowed to air dry overnight in a sealed laminar-flow hood. On the day of the assay, the residual PDL was removed, and the wells were washed with 90 μl of ddH2O. The water was then removed, and the plate was allowed to air-dry in the laminar-flow hood with a lid off. The bacteria were diluted to obtain 1 ml with an OD_600_ of 0.51 (this quantity of bacteria was determined to be within the reliable working range of the Seahorse XFP instrument after extensive optimization). These samples were then centrifuged for 10 minutes at 15,000 × g and room temperature to pellet the bacteria, and the supernatant was discarded. The bacteria were then resuspended in unbuffered 7H9 and centrifuged for 7 minutes at room temperature. The bacteria were resuspended in 1 ml of new unbuffered 7H9. A total of 90 μl of well-mixed bacterial solution was deposited in each well of the Seahorse XFP bacteria cell plate, and the plate was then centrifuged at 2,000 × g for 10 minutes. Subsequently, 90 μl of unbuffered 7H9 was gently added in a dropwise manner to increase the volume to 180 μl.

At least 2 hours after calibrant incubation in the utility plate-cartridge unit, the unit was ready for calibration in the Seahorse XFP instrument. Once calibrated, the utility plate was ejected and could be replaced with the bacterial cell plate. The instrument is returned to 37°C to begin the assay. Each measurement cycle requires 4 minutes of mixing and 2 minutes of sensor measurement. The assay consisted of three measurement cycles in the absence of a carbon source. Subsequently, 20 μl of 2% glucose-glycerol was injected to a final concentration of 0.2%, and 12 further measurements were obtained. The bacterial counts were normalized by CFUs of the different bacterial strains used to inoculate each well. The data were normalized to 10^5^ CFUs.

### Metabolite extraction experiments

For the targeted metabolomics profiling studies, the mycobacteria were initially grown in 7H9 liquid medium containing the carbon sources of interest until the OD_600_ reached approximately 0.8-1. The bacteria were then inoculated onto 0.22-μm nitrocellulose filters under vacuum filtration. The mycobacteria-laden filters were then placed on top of chemically equivalent agar media (described above) and allowed to grow at 37°C for 5 doubling times to generate sufficient biomass for targeted metabolomics studies. The filters were then transferred into 7H10 plates supplemented with 0.5 g/l Fraction V (bovine serum albumin), 0.2% dextrose, 0.2% glycerol, and 10 mM NaCl. The bacteria were metabolically quenched by plunging the filters into the extraction solution composed of acetonitrile/methanol/H_2_O (2:2:1) precooled to 4°C. Small molecules were extracted by mechanical lysis of the entire bacteria-containing solution with 0.1-mm acid-washed zirconia beads for 1 minute using a FastPrep (MP Bio®) set to 6.0 m/second. The lysates were filtered twice through 0.22-μm Spin-X column filters (CoStar®). The bacterial biomass of the individual samples was determined by measuring the residual protein content of the metabolite extracts using the BCA assay kit (Thermo®)^102,103^. A 100-μl aliquot of the metabolite solution was then mixed with 100 μl of acetonitrile with 0.2% acetic acid at −20°C and centrifuged for 10 minutes at 17,000 × g and 4°C. The final concentration of 70% acetonitrile was compatible with the starting conditions of HILIC chromatography. The supernatant was then transferred into LC/MS V-shaped vials (Agilent 5188-2788), and 4 μl was injected into the LC/MS instrument.

### Liquid chromatography-mass spectrometry for targeted metabolomics

Aqueous normal-phase liquid chromatography was performed using an Agilent 1290 Infinity II LC system equipped with a binary pump, temperature-controlled autosampler (set at 4°C) and temperature-controlled column compartment (set at 25°C) containing a Cogent Diamond Hydride Type C silica column (150 mm × 2.1 mm; dead volume of 315 μl). A flow rate of 0.4 mL/min was used. The elution of polar metabolites was performed using solvent A, which consisted of deionized water (resistivity ~18 MΩ cm) and 0.2% acetic acid, and solvent B, which consisted of 0.2% acetic acid in acetonitrile. The following gradient was used at a flow rate of 0.4 ml/min: 0 minute, 85% B; 0-2 minutes, 85% B; 3-5 minutes, 80% B; 6-7 minutes, 75% B; 8-9 minutes, 70% B; 10-11 minutes, 50% B; 11.1-14 minutes, 20% B; 14.1-25 minutes, 20% B; and 5-minute re-equilibration period at 85% B. Accurate mass spectrometry was performed using an Agilent Accurate Mass 6545 QTOF apparatus. Dynamic mass axis calibration was achieved by continuous infusion after chromatography of a reference mass solution using an isocratic pump connected to an ESI ionization source operated in the positive-ion mode. The nozzle and fragmentor voltages were set to 2,000 V and 100 V, respectively. The nebulizer pressure was set to 50 psig, and the nitrogen drying gas flow rate was set to 5 l/minute. The drying gas temperature was maintained at 300°C. The MS acquisition rate was 1.5 spectra/sec, and *m*/*z* data ranging from 50 to 1,200 were stored. This instrument enabled accurate mass spectral measurements with an error of less than 5 parts per million (ppm), a mass resolution ranging from 10,000 to 45,000 over the *m*/*z* range of 121-955 atomic mass units, and a 100,000-fold dynamic range with picomolar sensitivity. The data were collected in the centroid 4-GHz (extended dynamic range) mode. The detected *m*/*z* data were deemed to be metabolites identified based on unique accurate mass-retention time and MS/MS fragmentation identifiers for masses exhibiting the expected distribution of accompanying isotopomers. The typical variation in abundance for most of the metabolites remained between 5 and 10% under these experimental conditions.

### Liquid chromatography-mass spectrometry for untargeted metabolomics

The data were acquired with an Agilent 1290 Infinity II UHPLC coupled to a 6546 LC/Q-TOF system. Chromatographic separation was performed with an Agilent InfinityLab Poroshell 120 HILIC-Z (2.1 × 100 mm, 2.7 μm (p/n 675775-924)) column. The HILIC methodology was optimized for polar acidic metabolites. For easy and consistent mobile-phase preparation, a concentrated 10x solution consisting of 100 mM ammonium acetate (pH 9.0) in water was prepared to produce mobile phases A and B. Mobile phase A consisted of 10 mM ammonium acetate in water (pH 9) with 5-μm Agilent InfinityLab Deactivator Additive (p/n 5191-4506), and mobile phase B consisted of 1.0 mM ammonium acetate (pH 9) in 10:90 (v:v) water/acetonitrile. The following gradient was used with a flow rate of 0.5 ml/min: 0 minute, 100% B; 0-11.5 minutes, 70% B; 11.5-15 minutes, 100% B; 12-15 minutes, 100% B; and a 5-minute re-equilibration period at 100% B. Accurate mass spectrometry was performed using an Agilent Accurate Mass 6545 QTOF apparatus. Dynamic mass axis calibration was achieved by continuous infusion after chromatography of a reference mass solution using an isocratic pump connected to an ESI ionization source operated in the negative-ion mode. The following parameters were used: sheath gas temperature, 300°C; nebulizer pressure, 40 psig; sheath gas flow, 12 l min^−1^; capillary voltage, 3000 V; nozzle voltage, 0 V and fragmentor voltage, 115 V. The data were collected in the centroid 4 GHz (extended dynamic range) mode.

### cAMP standard curve

A stock solution of 3’,5’-cAMP at a concentration of 100 mM in double-distilled water was prepared and serially diluted in a solution of acetonitrile/methanol/H_2_O (2:2:1) to obtain concentrations in the range of 1 to 0.1 nM in technical quadruplicates. A standard curve was established using Agilent Quantitative Analysis B.07.00.

### Stable isotope labelling

Under the experimental conditions described above using [U-^13^C_3_]-glycerol (99%) and [U-^13^C_6_]-glucose (99%), the extent of ^13^C labelling of each metabolite was determined by dividing the summed peak height ion intensities of all ^13^C-labelled species by the ion intensity of both labelled and unlabelled species using Agilent Profinder version B.8.0.00 service pack 3.

### Statistical analysis

The data are presented as the means±SDs from at least two biological replicates and three technical replicates per condition. Unpaired two-tailed Student’s *t*-tests were used to compare the data, and p<0.05 was considered significant.

### Biological safety considerations

The bacteria were handled within a Class-II safety-level cabinet equipped with a UV light source and HEPA filters.

**Figure 7:**
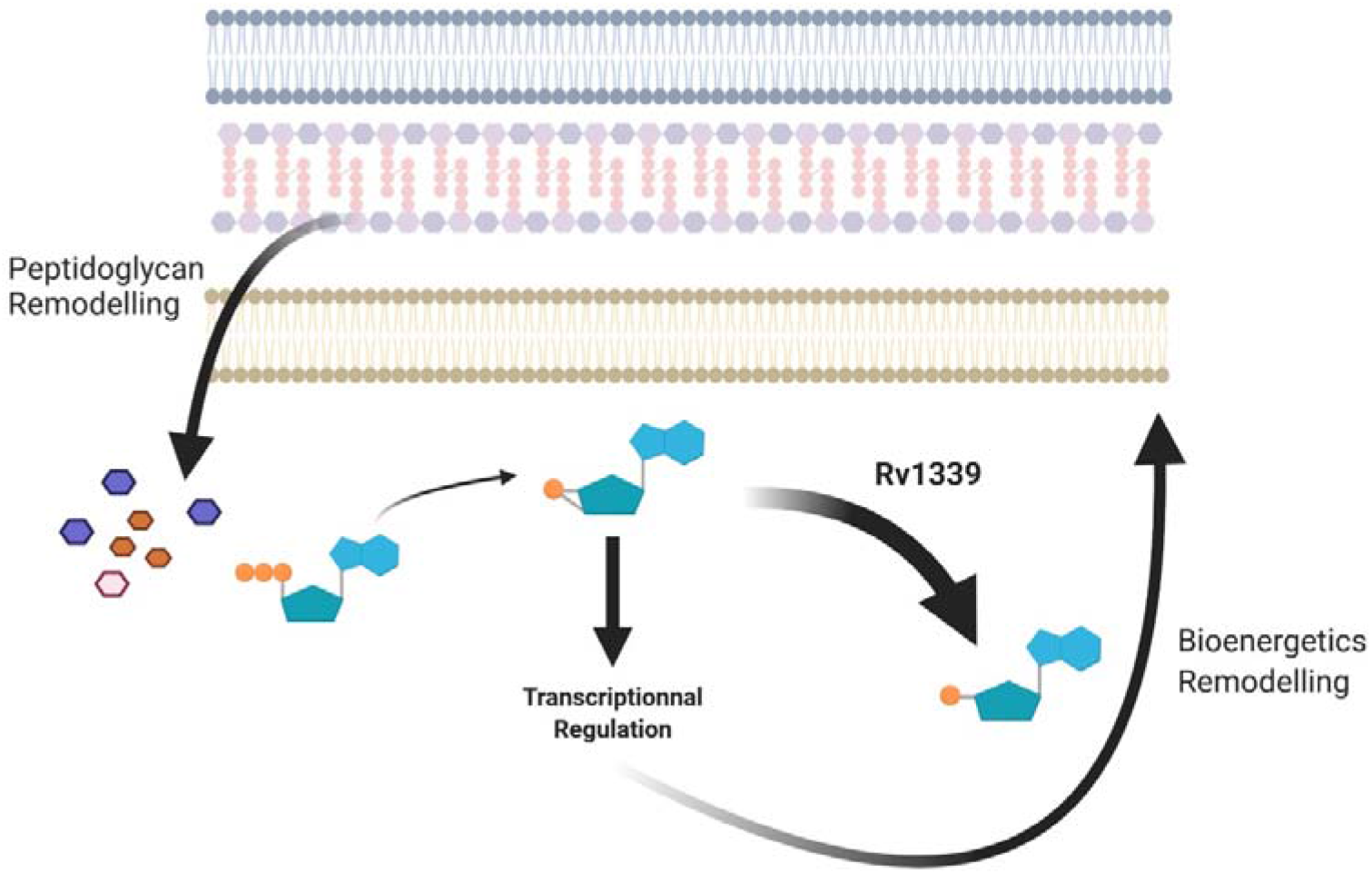
Model.

## Supporting information

Supplementary Informations

## Acknowledgements

We would like to thank Prof. Sandhya Visweswariah from the Indian Institute of Science, Bangalore, India, for providing the *M. bovis* BCG strains used in this work and Dr. Despoina Mavridou and Prof. Jose Penades for careful reading of the manuscript.

## Data and code availability

The RNA-seq data are available at GEO under the accession number GSE (pending) and the accompanying code is available at https://github.com/ash-omics/cAMP_RNAseq.

## Author contribution

MT, KN and GLM conceived and designed the experiments. MT, KN, YL, AC, AGG, and GLM performed the experiments, analysed the data, and wrote the paper.

## Additional information

The authors declare that they have no conflicts of interest regarding the content of this article.

