## Supplementary Informations for "Modulation of the cAMP levels with a conserved actinobacteria phosphodiesterase enzyme reduces antimicrobial tolerance in mycobacteria"

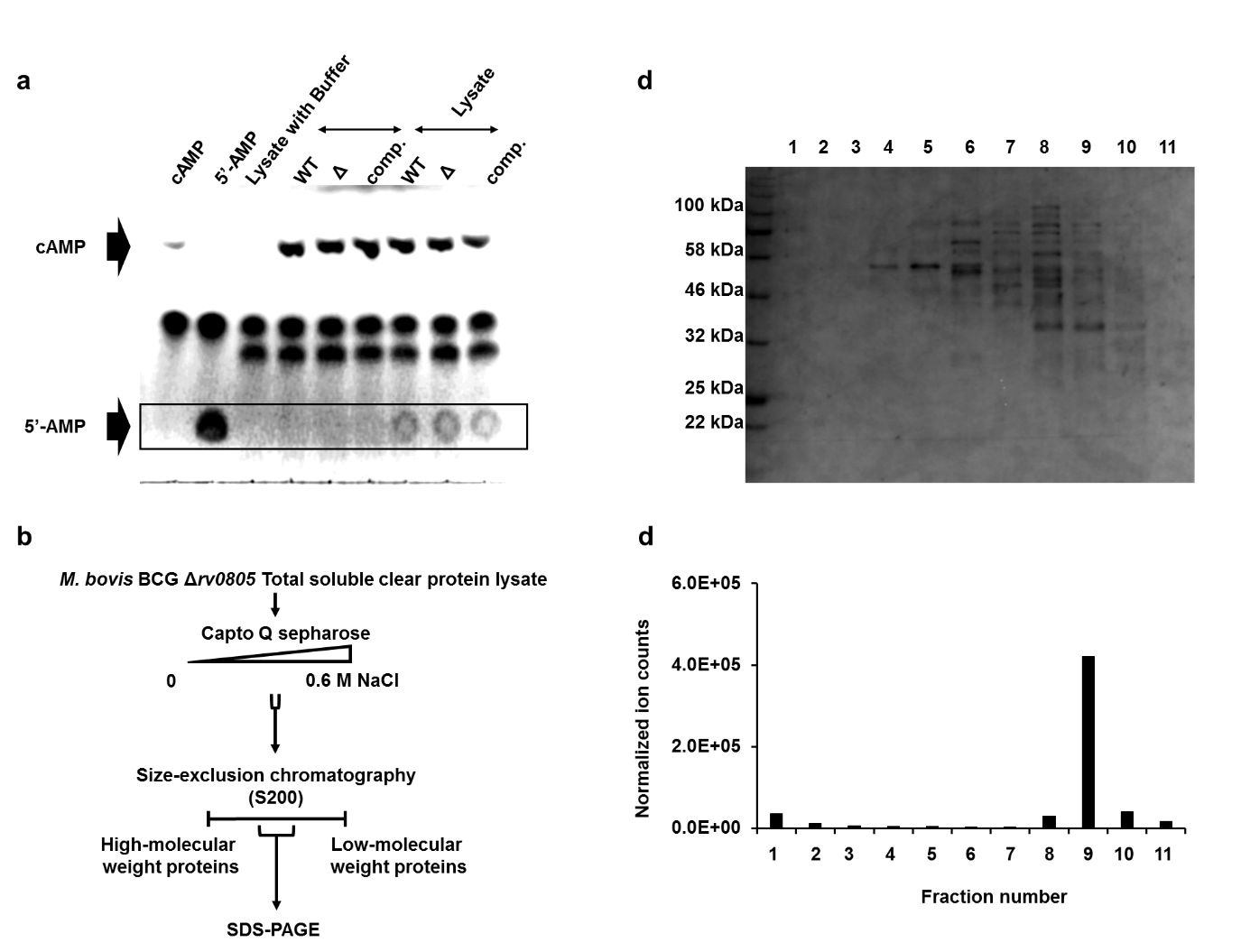

**Supplementary Figure 1: Identification and predicted 3-D structure of Rv1339**. **a**. TLC showing the *in vitro* cAMP activity of clear lysate from parental *M. bovis* BCG and strains with deletion (Δ) and complementation (comp.) of the *Rv0805* gene. **b**. Strategy for identifying the PDE activity of the Δ*rv0805* clear lysate using a series of liquid chromatography techniques. **c.** SDS-PAGE showing the protein content of the fractions collected from the last chromatography step. **d**. Bar graph displaying the ion counts normalized to the amount of 5’-AMP protein.

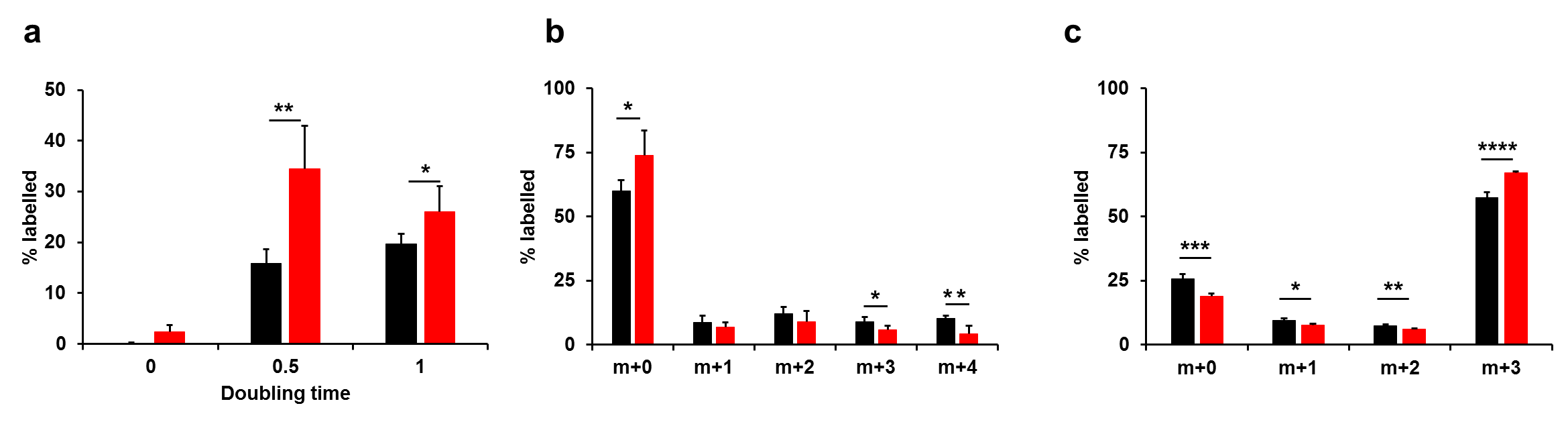

**Supplementary Figure 2: The expression of Rv1339 leads to changes in carbon turnover**. **a.** Percentage of labelled cAMP. **b.** Isotopologue distribution of succinate. **c.** Isotopologue distribution of serine. The data are presented as the means±SDs from two biological replicates and three replicates.

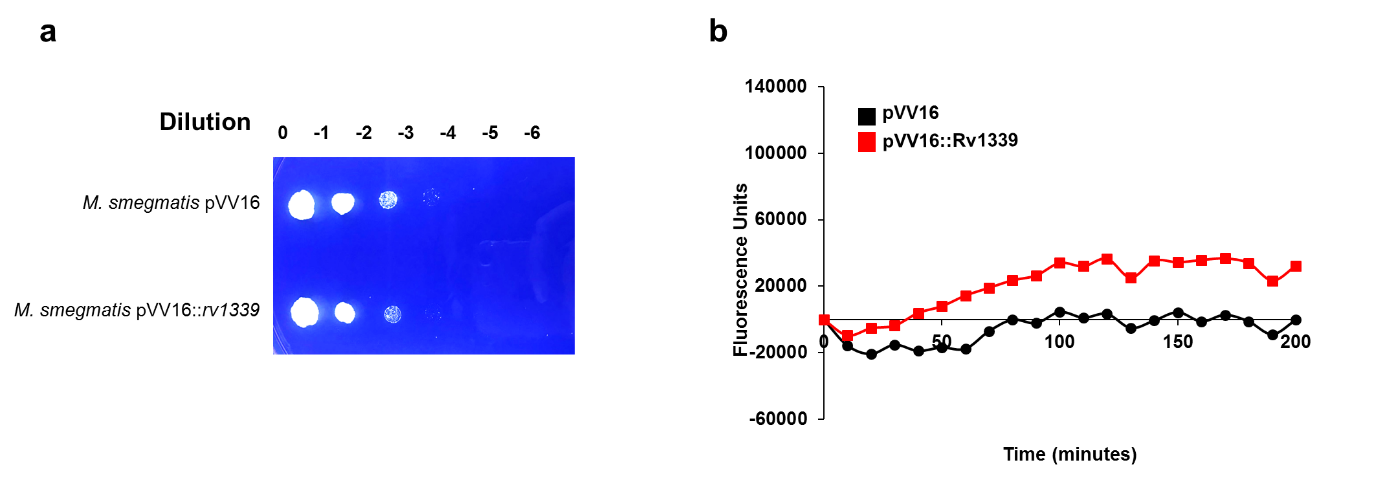

**Supplementary Figure 3:** **The expression of Rv1339 leads to an increase in the cell envelope permeability. a.** Crystal violet spot assay. **b.** Hoechst staining assay.

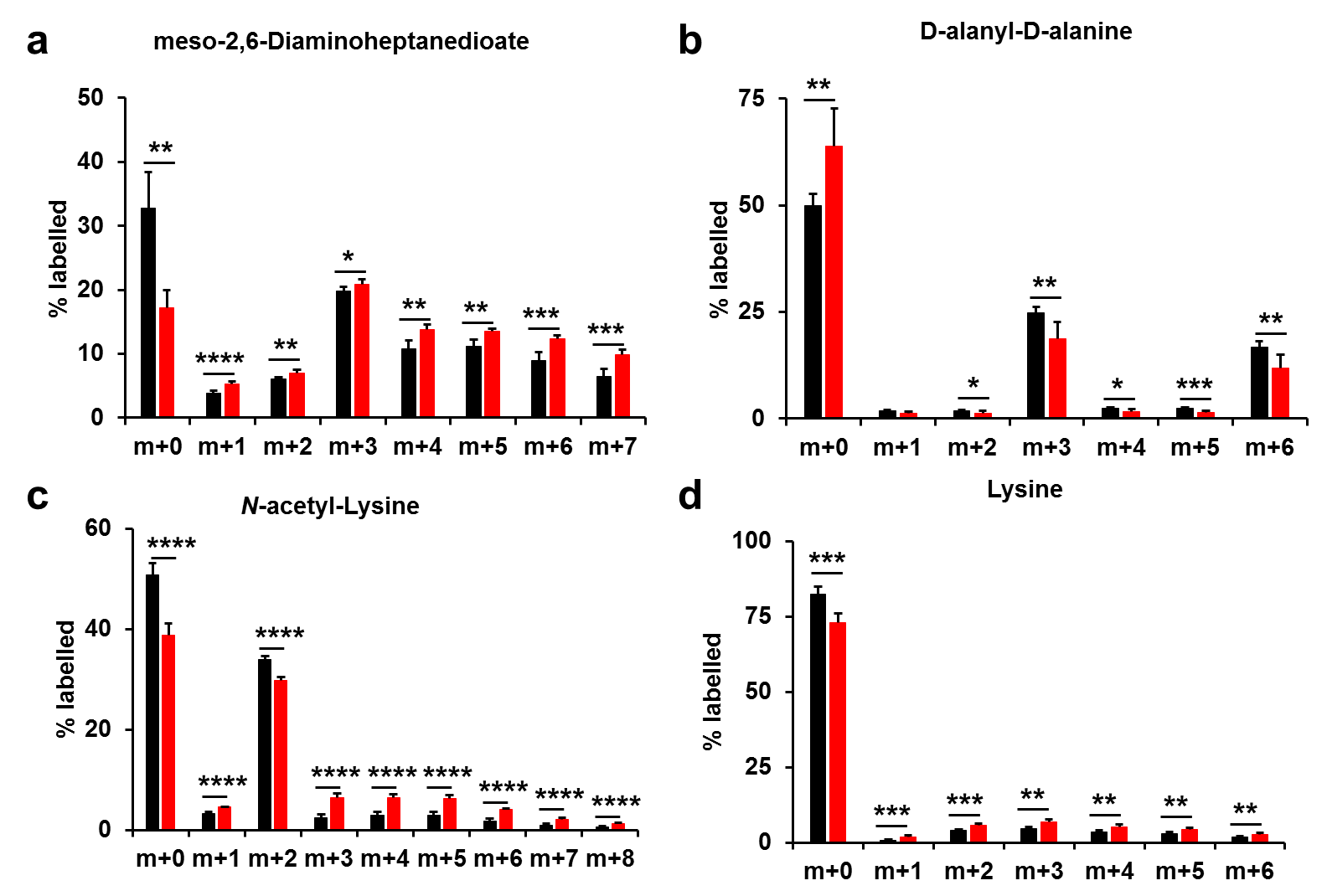

**Supplementary Figure 4:** **The expression of Rv1339 leads to an increase in the turnover of metabolites involved in peptidoglycan synthesis. a**. Isotopologue distribution of meso-2,6-diaminoheptanedioate. **b**. Isotopologue distribution of D-alanyl-D-alanine. **c**. Isotopologue distribution of N-acetyl-lysine. **d.** Isotopologue distribution of lysine. The data are presented as the means±SDs from two biological replicates and three replicates.

**Supplementary Table 1: Doubling time of the *M. smegmatis* mc^2^155 strains used in the study.**

| **Strains** | **Doubling time**  **(hours)** | ***p* value compared**  **with the empty vector-expressing strain** |
| --- | --- | --- |
| *M. smegmatis* pVV16 | 3.03 ± 0.07 | - |
| *M. smegmatis* pVV16::*rv1339* | 3.70 ± 0.21 | 0.001 |
| *M. smegmatis* pVV16::rv1339 D180A | 3.07 ± 0.09 | ns |

**Supplementary Table 2: MIC_90_ values of the *M. smegmatis* mc^2^155 strains used in the study.** The data are presented as the means±SDs from three biological replicates and three replicates.

| **Antibiotics** | **MIC (μg/ml) of**  ***M. smegmatis* pVV16** | **MIC (μg/ml) of**  ***M. smegmatis* pVV16::*r1339*** |
| --- | --- | --- |
| Ethambutol | 2.0 ± 0.0 | 2.0 ± 0.0 |
| Streptomycin | 0.50 ± 0.0 | 0.25 ± 0.0 |

**Supplementary Table 3: List of differentially expressed genes in Rv1339-expressing *M. smegmatis* mc^2^155.**

**Supplementary Table 4: List of differentially expressed metabolites in Rv1339-expressing *M. smegmatis* mc^2^155.**

| **Compound name** | **Chemical formula** | **Mass** | **Retention time (min)** | **Theoretical *m/z* (neg)** | **Experimental *m/z* (neg)** | **Δppm** |
| --- | --- | --- | --- | --- | --- | --- |
| Serine | C3H7NO3 | 105.042593 | 8.5 | 104.0353 | 104.03536 | -0.6 |
| Succinic acid | C4H6O4 | 118.026609 | 1.1 | 117.0193 | 117.01908 | 1.9 |
| Lysine | C6H14N2O2 | 146.1055 | 10 | 145.0983 | 145.09813 | 1.2 |
| *D*-Alanyl-D-alanine | C6H12N2O3 | 160.0848 | 5.6 | 159.0775 | 159.0774 | 0.6 |
| *L*-2-aminoadipic acid | C6H11NO4 | 161.0688 | 7.9 | 160.0615 | 160.0614 | 0.6 |
| *L*-2,3,4,5-Tetrahydrodipicolinate | C7H9NO4 | 171.0532 | 5 | 170.0459 | 170.04425 | 9.7 |
| meso-2,6-Diaminoheptanedioate | C7H14N2O4 | 190.095357 | 9.2 | 189.08808 | 189.08749 | 3.1 |
| *N*-Acetyl-alpha-*D*-glucosamine 1-phosphate | C8H16NO9P | 301.0563 | 8.4 | 300.049 | 300.04911 | -0.4 |

**Supplementary Table 5: Percentage of labelled metabolites altered in Rv1339-expressing *M. smegmatis* mc^2^155.** The data are presented as the means±SDs from two biological replicates and three replicates.

|  |  | **Percentage of labelled metabolites** | |  |
| --- | --- | --- | --- | --- |
| **Pathways** | **Metabolites** | ***M. smegmati*s pVV16** | ***M. smegmatis* pVV16::rv1339** | **p value** |
| ETC | succinate | 40.1 ± 4.2 | 26.1 ± 9.5 | 0.0199 |
| Glycolysis | serine | 74.2 ± 1.9 | 80.9 ± 0.9 | 0.0015 |
| Peptidoglycan remodelling | meso-2,6-diaminoheptanedioate | 66.3 ± 5.8 | 83.9 ± 0.7 | 0.0004 |
|  | lysine | 17.6 ± 2.2 | 26.9 ± 3.4 | 0.0003 |
|  | *D*-alanyl-*D*-alanine | 50.3 ± 3.1 | 38.3 ± 5.6 | 0.0031 |
|  | *N*-acetyl-lysine | 48.9 ± 2.5 | 61.9 ± 2.0 | 4E-05 |
